# Tensor-based representation learning for multi-omics integrative clustering and feature discovery

**DOI:** 10.64898/2025.11.28.691099

**Authors:** Yang Zhang, Lin Liu, Zhaoqi Liu, Qi Liu, Lina Ma, Zhang Zhang

**Author notes:** Beijing Forestry University, Beijing 100083, China (Lin Liu).

## Abstract

Multi-omics integrative analysis provides a powerful means for elucidating complex molecular mechanisms and biological processes, yet remains challenging in effectively representing the multi-dimensional relationships inherent to multi-omics data. Here we present MIA, a tensor-based representation learning framework that preserves the multi-dimensional structure of multi-omics data for accurate sample clustering and feature discovery. Unlike existing algorithms that primarily rely on two-dimensional representations, MIA models multi-omics data as a three-dimensional tensor and integrates tensor decomposition, fuzzy c-means, and an enhanced random forest model within a unified framework for clustering and feature discovery. Benchmarking on simulated and empirical datasets demonstrates that MIA consistently outperforms representative state-of-the-art algorithms in both clustering and feature identification. Application to multiple TCGA cancer types further shows its ability to stratify samples and identify molecular features associated with clinically relevant outcomes. Specifically, in glioblastoma, MIA reveals three previously uncharacterized subtypes with distinct prognostic profiles and uncovers feature genes strongly associated with subtype identity. These genes are further linked to therapeutic response and retain discriminative power across major glioblastoma cellular populations at single-cell resolution. Collectively, our results establish MIA as a generalizable computational framework for multi-omics integrative analysis, enabling systematic molecular stratification and interpretable feature discovery across diverse biological systems.

## Introduction

Powered by high-throughput sequencing technologies, data from multiple omics layers, such as genomics, transcriptomics, proteomics, and epigenomics, have been generated at exponential rates and scales^1^. Integrative analysis of multi-omics data can capture complementary information from their underlying relationships and thus holds great promise for elucidating complex molecular mechanisms and biological processes^2, 3^, particularly in the context of precision medicine^4^, through precise sample clustering^5–7^ and accurate identification of feature genes as potential molecular biomarkers^8, 9^. In machine learning and artificial intelligence, a feature is an individual characteristic of a data set and selection of informative and discriminative features is crucial for effective sample clustering. Hence, in biomedical research, a feature gene reflects a specific biological process and a set of feature genes is defined as those that contribute significantly to clustering. Therefore, multi-omics integrative analysis represents a powerful approach for systematically delineating feature genes from diverse omics data and thereby enabling a comprehensive understanding of complex physiopathological processes and mechanisms^10, 11^.

Over the past several years, several algorithms have been developed for multi-omics data integrative analysis, which roughly fall into two groups in light of data integration strategy^12, 13^: similarity network and consensus factor. For the former, representative algorithms include SNF^14^, PINS^15^, NEMO^16^, and MANAclust^17^. They calculate sample similarities and fuse networks across different omics types, but are sensitive to data perturbations, outliers and noise, which can reduce robustness and increase computational complexity. For the latter, representative algorithms are MOFA^18^, iClusterPlus^19^, and intNMF^20^. They derive shared latent factors across multi-omics types but often rely on specific statistical assumptions that may not apply universally across diverse biological datasets, thus rendering them susceptible to model misspecification and noise. Importantly, both strategies transform multi-omics data into two-dimensional representations, which may constrain the preservation of complex relationships across molecular layers and limit the joint characterization of complementary information. Among these existing algorithms, only SNF, MANAclust, and iClusterPlus are capable for feature gene identification as well as clustering. Yet, these three algorithms identify feature genes independently within each omics layer, rather than jointly across multi-omics layers.

A promising strategy to overcome these limitations is tensor-based representation learning, which provides a powerful framework to effectively capture high-dimensional features inherent in multi-omics data^21^. Specifically, tensor-based representation learning allows multi-omics data to be represented as multi-dimensional arrays, enabling the simultaneous encapsulation of sample, gene and omics type, without any prior assumption^22, 23^. In addition, the tensor framework exhibits unique advantages in handling high-dimensional heterogeneity of biomedical data, flexibly adapting to variable sample sizes ranging from dozens to several thousands, and accommodating ultra-high feature dimensionality greater than 10,000 features per omics type^24^, as testified by successful applications in sample classification^25, 26^, survival prediction^27^, and gene interaction prediction^28^. Despite these advances, tensor-based representation learning has not yet been applied to multi-omics clustering and feature discovery.

Toward this end, here we present MIA, a tensor-based representation learning algorithm for integrative multi-omics clustering and feature discovery. Unlike extant algorithms that rely on the two-dimensional data representation, MIA preserves the multi-dimensional structure of multi-omics data by modeling them as a three-dimensional tensor and learning compact representations through tensor decomposition. It further uses fuzzy c-means (FCM) for clustering to accurately capture sample similarity while minimizing the influence of initial cluster centers. For feature gene identification, we develop a new enhanced random forest model (ERFM) for the tensor representation, enabling feature contributions to be evaluated jointly across multi-omics layers rather than independently within each omics layer. To assess the power of MIA, we conduct extensive comparative analyses using both simulated and empirical datasets and apply it to glioblastoma (GBM) subtyping and feature gene identification.

## Results

### MIA framework

We modularize the framework of MIA into three modules, namely, multi-omics integration, tensor decomposition & clustering, and feature gene identification (**Fig. 1**). For multi-omics integration, MIA represents multi-omics data as a three-dimensional tensor comprising samples, genes, and omics types (**Fig. 1a**). It preserves the structural relationships within the data and facilitates the learning of joint representations across multi-omics types.

**Fig. 1.**
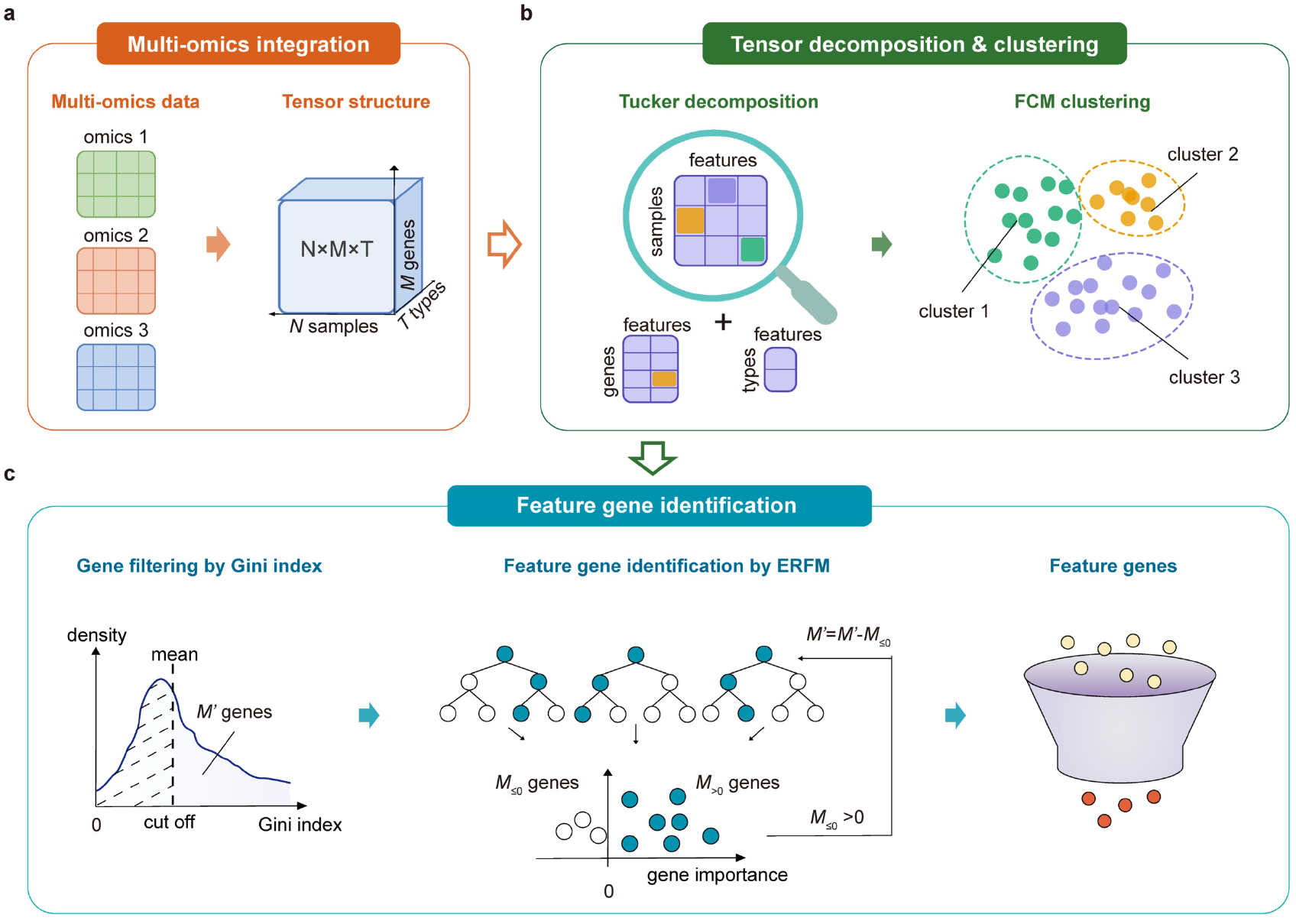
Schematic of the MIA framework. (a) Various types of omics data are integrated into a three-dimensional tensor comprising samples, genes, and omics types. (b) Tucker decomposition is used to decompose the tensor into three two-dimensional factor matrices: sample-feature matrix, gene-feature matrix, and type-feature matrix. Subsequently, FCM clustering is applied to the sample-feature matrix. (c) Feature gene identification is performed based on the tensor and sample labels or clustering results. After gene filtering by the Gini index, an ERFM designed for three-dimenstional tensor is applied to quantify the contribution of each gene in clustering and thus to yield a set of feature genes (see details in Methods).

To analyze unlabeled biomedical samples, MIA applies tensor decomposition to learn a compact representation of the integrated tensor, followed by FCM clustering for sample stratification. Among available tensor decomposition techniques, Tucker decomposition is widely adopted, owing to its flexibility and accuracy in capturing complex data structures and interdependencies^21, 23^. Therefore, MIA adopts Tucker decomposition to factorize the tensor into three matrices: sample-feature matrix, gene-feature matrix, and omics type-feature matrix, and then employs FCM for clustering based on the sample-feature matrix (**Fig. 1b** and **Supplementary Fig. 1**). Unlike *K*-means, FCM allows samples to be classified into multiple clusters, providing more accurate characterization of ambiguous samples. As noted, there are two critical parameters for FCM—uncertain fuzzy index (*m*) and cluster count (*K*)—that are typically pre-defined. In MIA, these two parameters are optimally determined for each dataset: *m* is estimated from sample and feature counts^29^ and *K* is estimated by maximizing the silhouette coefficient, thereby providing data-driven parameter estimation tailored to each dataset (see details in Methods; **Supplementary Figs. 2 and 3**).

Using the clustering results as sample labels, MIA first utilizes the Gini index to perform preliminary gene filtering and then employs ERFM to identify feature genes based on their joint contributions across omics layers. Unlike traditional random forest model (RFM) that handles only a single data type, ERFM accounts for multi-omics layers when quantifying the contribution of each gene to sample clustering and thus to yield a set of feature genes (**Fig. 1c**). There are two significant improvements in ERFM: an enhanced tree structure and an iterative computational procedure (**Supplementary Fig. 4**). First, the enhanced tree structure accounts for multi-dimensional data types by creating multiple tree nodes based on different omics types for each gene, rather than a single tree node in RFM. The importance score of each gene is determined by the increase of classification error after simultaneous randomizing its corresponding nodes across omics layers, allowing gene contributions to be assessed jointly rather than independently within individual omics layers. Second, instead of a single forest generation in RFM, ERFM employs an iterative computational procedure that repeatedly constructs and refines the forest, thus enhancing the consistency of the identified feature gene set. Specifically, after each iteration, genes with negative importance scores are eliminated, and the iterative refinement continues until there is no remaining gene with negative importance scores, ensuring that only the most relevant feature genes are retained for further analysis (**Supplementary Fig. 4c,d**; see details in Methods).

### Benchmarking on simulated datasets

To evaluate the performance of MIA, we consider multiple parameters, including cluster count (*K*), omics type count (*T*), sample count (*N*), and the numbers of real (true) features (*M_t_*) and noise (false) features (*M_f_*), and generate simulated datasets with different parameter combinations in terms of possible scenarios of real data (see details in Methods). We examine the clustering accuracy of MIA by comparison against six state-of-the-art algorithms—SNF, MANAclust, iClusterPlus, PINS, NEMO and MOFA—among which only the former three support feature gene identification.

When varying *K* and *T* with fixed *N*, *M_t_* and *M_f_*, MIA overall achieves stable and high accuracy across all parameter combinations (**Fig. 2a**). Among these competing algorithms, NEMO and SNF present comparable performance, while iClusterPlus exhibits marked instability when varying *K.* Consistently, MIA outperforms these algorithms at varied values of *N, M_t_*, *M_f_* with fixed *K* and *T* (**Fig. 2b**). In contrast, MANAclust shows significant deviations from the true labels. Considering the significance of *K* in data analysis (that is often arbitrarily set), we further evaluate their accuracy in optimal *K* selection and reveal that MIA achieves an average accuracy of 0.987 across all datasets, outperforming these existing algorithms (**Fig. 2c**). Since MANAclust often overestimates the value of *K*, we exclude it from the following analysis in real datasets. Regarding feature gene identification, it is clearly shown that feature genes identified by MIA yield higher significance than those identified by other algorithms (**Fig. 2d and Supplementary Table 2**).

**Fig. 2.**
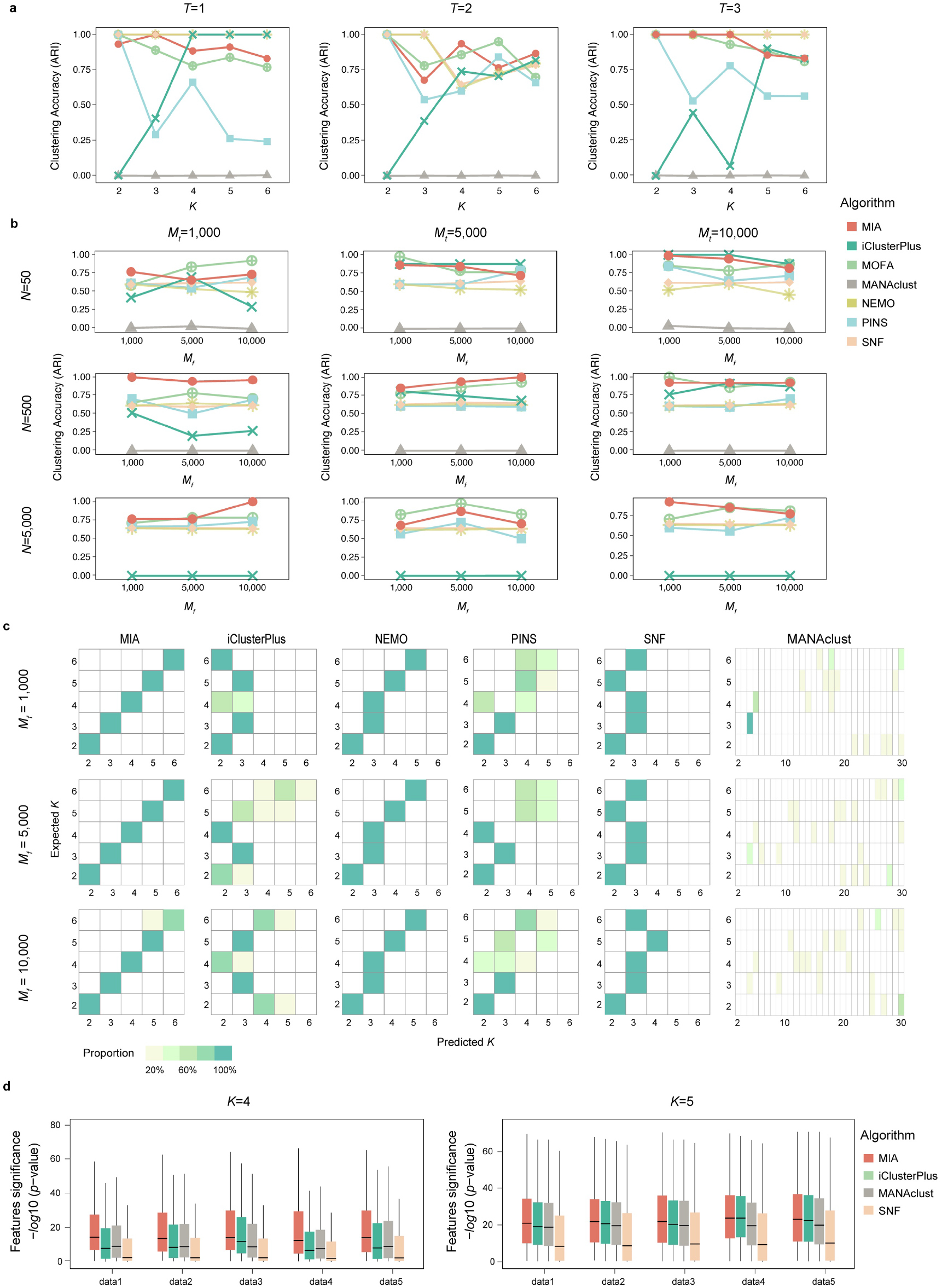
Performance of MIA in clustering and feature identification on simulated datasets by comparison against several existing algorithms. (a) Clustering performance across different numbers of clusters (*K*=2∼6) and omics types (*T*=1∼3), with fixed sample size (*N*=500), real features (*Mt*=5,000), and noise features (*Mf*=5,000). Clustering accuracy is assessed by the Adjusted Rand Index (ARI). (b) Clustering performance across varied sample sizes (*N*=50, 500, and 5,000), real features (*Mt*=1,000, 5000, and 10,000), and noise features (*Mf*=1,000, 5,000, and 10,000), with fixed *K*=4 and *T*=2. (c) Evaluation of optimal *K* estimation by different algorithms based on simulated datasets consisting of *K*=2∼6, *Mf*=1,000, 5,000, and 10,000, with fixed *N*=500 and *Mt*=1,000. The proportion of predicted *K* values identified by each algorithm is calculated across simulated replicates. (d) Performance of feature genes identified by different algorithms is evaluated by the Kruskal-Wallis test on five simulated replicates (data1∼data5) with *K*=4∼5, *T*=2, *N*=500, *Mt*=1,000, and *Mf*=10,000.

### Benchmarking on TCGA datasets

We evaluate MIA in ten empirical datasets from The Cancer Genome Atlas (TCGA), covering different omics data types (copy number variation (CNV), gene expression, and methylation) and involving ten cancer types: Bladder Cancer (BLCA), Cervical Squamous Cell Carcinoma (CESC), Breast Cancer (BRCA), Colon Cancer (COAD), GBM, Head and Neck Squamous Cell Carcinoma (HNSC), Lower Grade Glioma (LGG), Liver Cancer (LIHC), Lung Adenocarcinoma (LUAD), and Prostate Cancer (PRAD). To facilitate the comparison, we run MIA as well as the six algorithms to perform clustering and survival analyses on each dataset.

In these ten datasets, MIA overall outperforms these existing algorithms by achieving more significant survival differences across different omics data combinations (**Fig. 3a-c** and **Supplementary Table 3**), particularly for COAD and LUAD (**Supplementary Figs. 5 and 6**). Compared to all possible combinations of single-omics and two-omics data, clustering by MIA on three-omics data generally demonstrates higher survival significance (*p*<0.05, Log-rank test) in nearly all cancer types, which is more striking than that of other algorithms. Particularly, MIA achieves significantly different survival outcomes (*p*<0.0001, Log-rank test) in LIHC and GBM based on three-omics data, whereas single-omics or two-omics combinations fail to achieve similar differences (**Supplementary Figs. 7 and 8**). Additionally, we observe that although nearly all algorithms exhibit comparable survival differences in LGG on multi-omics combinations, clusters determined by MIA show more divergences (**Supplementary Fig. 9**). Detailed survival analyses across all these ten cancer types (**Supplementary Figs. 5-14**) show that multi-omics data integrative analysis by MIA is superior to other algorithms in precise clustering.

**Fig. 3.**
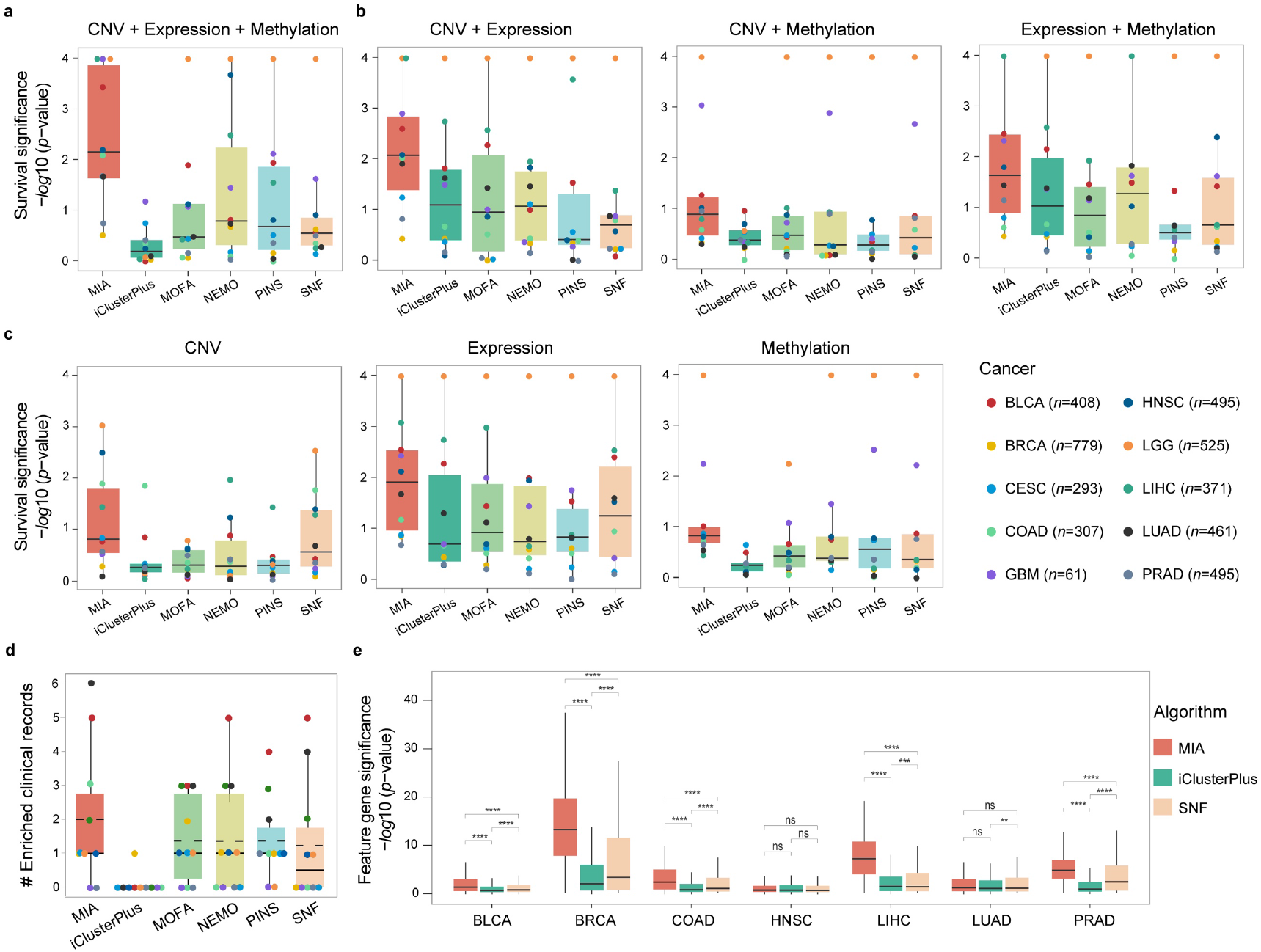
Evaluation of survival outcomes and feature genes determined by MIA on TCGA datasets in comparison with existing algorithms. Comparison of clustering results from different algorithms based on survival significance in (a) three-omics, (b) two-omics, and (c) single-omics datasets. (d) Number of significantly enriched clinical records (*p*<0.05) across ten cancer types. Horizontal lines indicate the median and dashed lines represent the mean. (e) Feature gene significance between cancer and normal samples across different cancer types. The Kruskal-Wallis test was used to evaluate the average significance of feature genes identified by each algorithm (*p*-value, with ****<0.0001, ***<0.001, **<0.01, *<0.05, and ns not significant).

Following a previous study^12^, we adopt six clinical records to evaluate the performance of MIA, namely, gender, age at diagnosis, tumor progression (pathologic T), metastases status (pathologic M), lymph node involvement (pathologic N), and overall pathologic stage, which can be used to indicate demographic characteristics, disease severity and progression. We investigate the enrichment of these clinical records in all clusters determined by different algorithms and find that MIA clustering achieves more significant clinical outcomes across ten cancer types (**Fig. 3d**, **Supplementary Fig. 15** and **Supplementary Table 5**). In BLCA, all the algorithms reveal more enriched records in a consistent manner, possibly due to its distinct clinical stages and molecular differences between non-muscle invasive and muscle-invasive types^30^. Notably, no clinical records are enriched in GBM across all algorithms, most likely owing to the missing clinical data as well as high malignancy of GBM that is difficult to distinguish subtypes based on tumor morphology and pathology. Given the significance of these six clinical records, our results show that MIA clustering has a stronger correlation with disease progression than other algorithms.

We further assess the MIA performance of feature gene identification in seven cancer types with sufficient samples. Compared with existing algorithms, feature genes identified by MIA present greater significant differences between cancer and normal samples across nearly all examined cancer types (**Fig. 3e**). Clearly, this result demonstrates the superior capability of MIA in feature gene identification from multi-omics integrative analysis.

### GBM subtyping and feature gene identification

GBM is the most aggressive primary brain cancer, with a five-year survival rate of only ∼5%^31^. Therefore, it is crucial to identify feature genes that can be used as potential biomarkers for GBM diagnosis and prognosis. We apply MIA to the GBM dataset described above, covering 15,372 genes with multi-omics profiles (CNV, gene expression and methylation). For clustering, MIA determines the optimal value of *K* at 3 (**Fig. 4a**), as further supported by the well-separated clusters (**Fig. 4b**). Most important, clinically, the clustering of *K*=3 yields more significant survival differences (*p*-value<0.0001, Log-rank test; **Fig. 4c**), namely, clusters 1, 2 and 3 at high, medium and low survivals, respectively. In contrast, classical subtypes based solely on gene expression^32^ show no significant survival differences (*p*-value=0.59, Log-rank test; **Supplementary Fig. 16**). Therefore, our results show that MIA is more powerful in multi-omics integrative clustering.

**Fig. 4.**
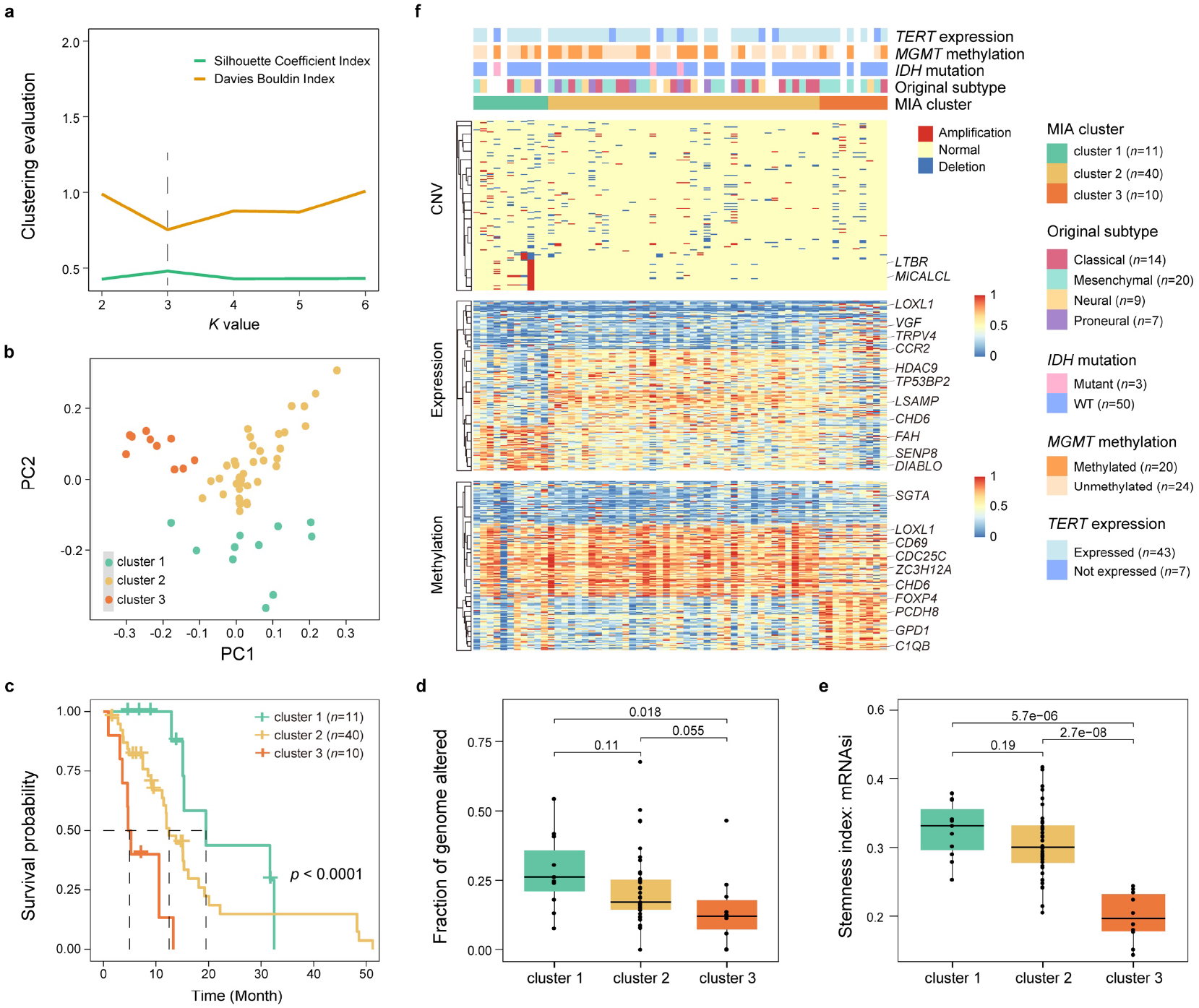
GBM subtyping analysis by MIA. (a) Clustering evaluation at different *K* values using the silhouette coefficient and Davies-Bouldin index. A higher silhouette coefficient and lower Davies-Bouldin index indicate better cluster separation. (b) Principal component analysis (PCA) of the sample-feature matrix obtained from Tucker decomposition. (c) Kaplan-Meier survival curves for the three clusters identified by MIA. The log-rank test was used to assess the statistical significance. (d) Fractions of genome altered across the three clusters. (e) Stemness index (mRNA expression-based stemness index, mRNAsi) across the three clusters, with statistical significance assessed by the Wilcoxon rank-sum test. (f) Heatmap of 167 GBM feature genes identified by MIA.

It has been reported that the fraction of altered genome reflects the extent of genetic alterations in cancer^33^ and stemness index signifies the stem cell-like characteristics of cancer cells^34^, together indicating tumor aggressiveness and prognosis. Thus, we further evaluate the MIA clustering in GBM based on these metrics. We find that the fraction of altered genome decreases significantly from cluster 1 to cluster 3 (*p*<0.05; **Fig. 4d**), corresponding well with the declining survival rates. Consistently, the stemness index gradually is on the decrease, namely, higher in cluster 1, medium in cluster 2, and lower in cluster 3 (**Fig. 4e** and **Supplementary Table 6**). These results indicate that GBM with fewer genomic alterations may exhibit more aggressive behavior and lower survival rates, consistent with observations in advanced tumors^33^. In addition, GBM with poor prognosis has lower stemness and higher cancer cell differentiation^35, 36^. Overall, the clinical analysis demonstrates that MIA is effective to identify GBM subtypes with distinct aggressiveness and prognosis.

In addition, MIA is capable to identify a set of 167 feature genes from the GBM three-omics profiles that contribute to the subtyping (**Fig. 4f and Supplementary Table 7**). Among them, 87 (52.1%) genes have already been documented to be associated with GBM, in which 70 are recognized as GBM markers (**Fig. 5a**). Clusters 1, 2, and 3, built on these 167 feature genes, exhibit distinct molecular characteristics, with representative reported markers as listed below; cluster 1 shows copy number amplification in *LTBR* and *MICALCL* and increased expression in *FAH*, *SENP8*, and *DIABLO*, cluster 2 exhibits increased expression in *HDAC9*, *LSAMP*, and *CHD6*, and cluster 3 presents increased expression in *VGF*, *TRPV4*, and *CCR2* and increased methylation in *PCDH8*, *GPD1*, and *C1QB* (**Fig. 4f**). Furthermore, tissue enrichment and functional annotation analyses reveal that the 167 feature genes are closely associated with brain tissues and crucial for pathobiological processes, such as neuroblast proliferation, immune microenvironment dynamics, and tumor-related pathways (**Supplementary Fig. 17a,b**).

**Fig. 5.**
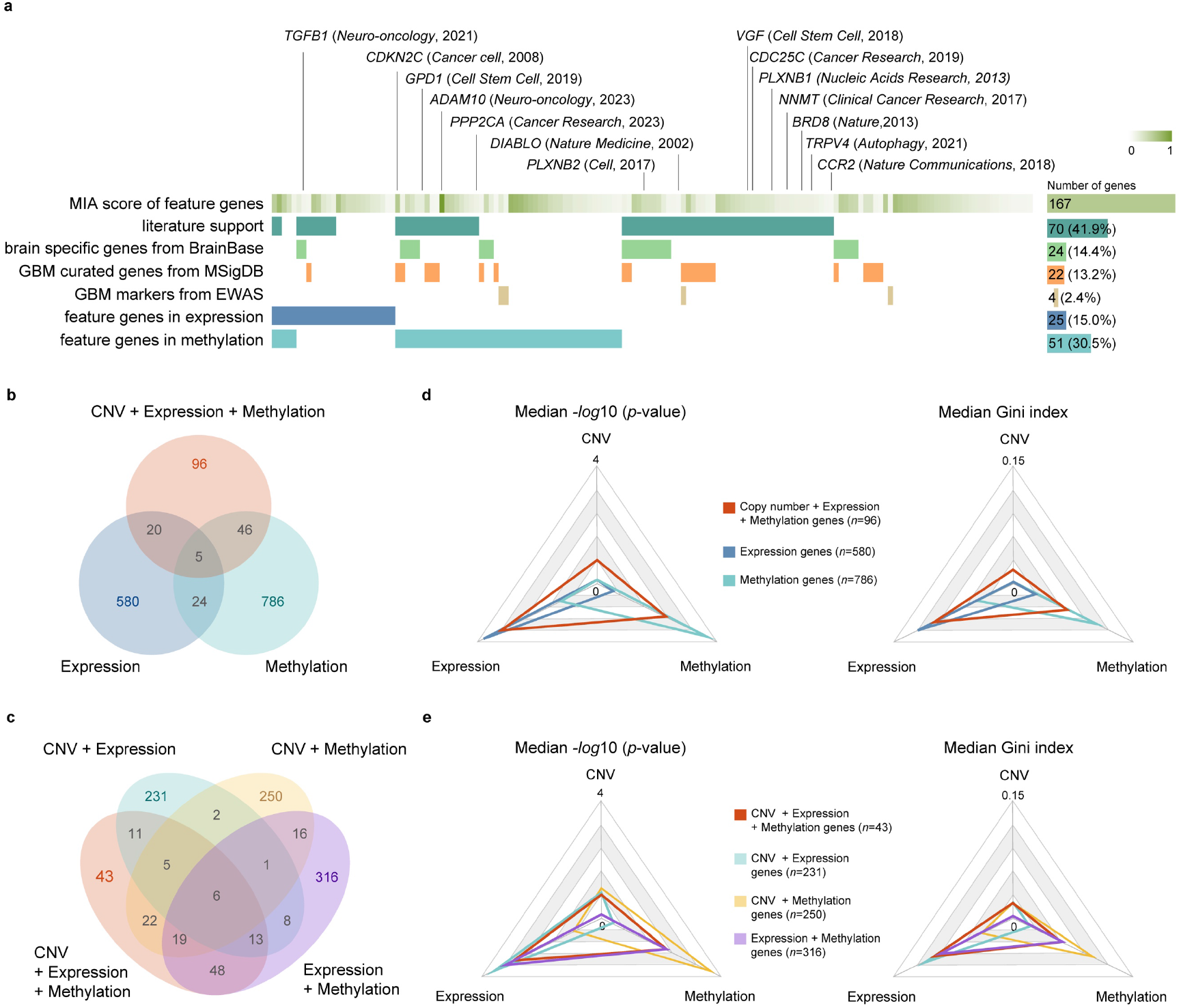
Characterization of 167 GBM feature genes identified by MIA. (a) Feature genes ranked by MIA scores (0∼1), with literature evidence, database annotations, and genes identified by single-omics data analyses (genes identified by CNV did not reach the threshold) (**Supplementary Table 9**). (b-c) Feature genes identified from three-omics, by comparison with single-omics and two-omics, respectively. (d-e) Clustering power of feature genes specifically identified from three-omics, by comparison with single-omics and two-omics, respectively. The Kruskal-Wallis test and the Gini index were employed to evaluate the clustering power.

We also use MIA to identify feature genes based on single-omics and two-omics data. Among these 167 feature genes, 96 (57.5%) and 43 (25.7%) genes are specifically identified based on three-omics data, by comparison against single-omics and two-omics data, respectively (**Fig. 5b,c**). Functional enrichment analysis shows that these feature genes specifically identified by three-omics data exhibit stronger associations with brain tissue and tumor development (**Supplementary Fig. 17c**). We further analyze the clustering power of feature genes specifically identified by different omics combinations. We observe that feature genes identified from single-omics data exhibit strong significance within their respective omics layers, but feature genes from three-omics always show higher clustering scores, clearly indicating that multi-omics integrative analysis achieves more enhanced clustering power (**Fig. 5d,e**). Notably, several literature-reported GBM marker genes, such as *VGF* (promoting GBM stem cell survival and tumor growth^37^) and *BRD8* (hindering *p53* transcriptional activation and promoting GBM cell proliferation^38^), are successfully identified by three-omics data integrative analysis, yet absent in single-omics or two-omics data analysis (**Supplementary Table 8**). It is worth mentioning that based on three-omics integrative analysis, *LTBR* (activating anti-tumor immune response in glioma^39^) and *GPD1* (whose inhibition prolongs GBM mouse survival^40^) are computationally identified as feature genes with markedly enhanced priority (**Supplementary Fig. 18**), thus highlighting the great potential of MIA to uncover biologically significant features from multi-omics data and thereby guiding in-depth experimental validation and mechanism investigation.

### Characterization of Temozolomide treatment sensitivity through GBM feature genes

Temozolomide (TMZ) is one of the most effective chemotherapy agents for GBM treatment^30^. Based on these 167 feature genes identified by MIA, we further explore whether these genes can be used to predict TMZ chemotherapy response. Toward this end, firstly, we collect a publicly available dataset (expression data only) with GBM TMZ treatment, containing samples of control, TMZ + radiotherapy, and TMZ + radiotherapy + inhibitor^41^. MIA clustering based on the 167 gene expression profiles separates the samples into three clusters that correspond closely to the three treatment groups. We find that the control samples are most enriched in cluster 1, the TMZ + radiotherapy samples are most in cluster 2, and the TMZ + radiotherapy + inhibitor samples are exclusively included in cluster 3 (**Fig. 6a** and **Supplementary Table 9**). A multi-class ROC analysis yields an average AUC of 0.85, indicating that these genes effectively distinguish different TMZ treatment regimens (**Fig. 6b**). To further assess the relevance of these genes to TMZ treatment sensitivity, we collect three independent datasets containing TMZ-resistant and TMZ-sensitive GBM cell-line samples. For these three datasets, clustering results align perfectly with the outcomes of TMZ treatment (**Fig. 6c-e** and **Supplementary Table 9**), implying that these 167 genes are closely associated with TMZ chemotherapy response.

**Fig. 6.**
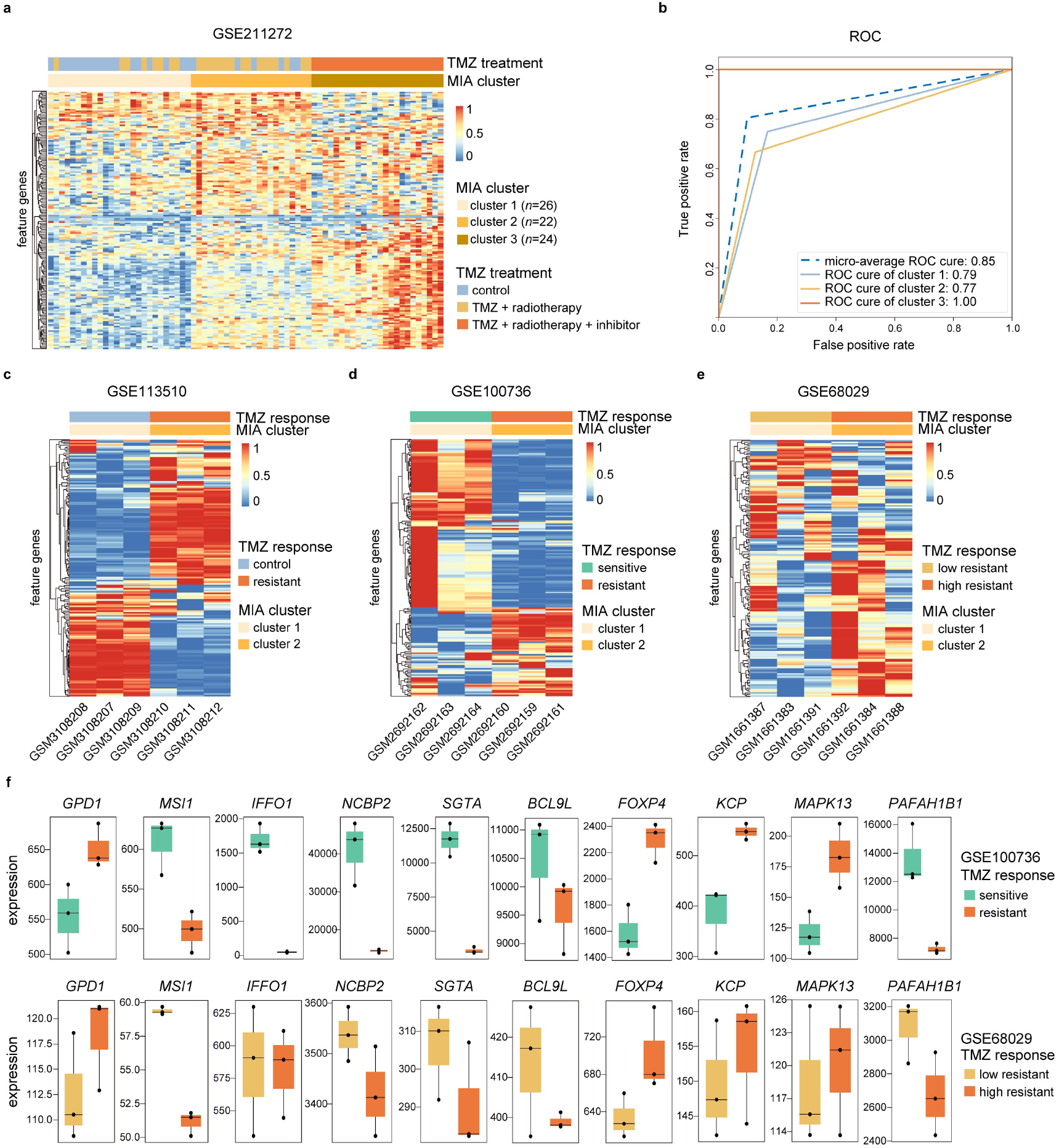
MIA clustering of 167 GBM feature genes in TMZ treatment datasets. (a) MIA clustering of 167 GBM feature genes in GSE211272, including control, TMZ + radiotherapy, and TMZ + radiotherapy + inhibitor samples. (b) Multi-class ROC curve illustrating clustering performance on GSE211272. (c-e) MIA Clustering of 167 GBM feature genes in three TMZ treatment datasets, including GSE113510 (control vs resistant), GSE100736 (resistant vs sensitive), and GSE68029 (low resistant vs high resistant). (f) The literature-supported feature genes, exhibiting consistent expression patterns in both GSE100736 and GSE68029.

To systematically identify genes associated with TMZ chemotherapy sensitivity, we discover that 64 genes display consistent expression patterns in the above datasets. We collect these 64 genes’ associations with chemotherapy drugs across different cancers (**Supplementary Table 10)**. Notably, ten genes (*GPD1*, *MSI1*, *IFFO1*, *NCBP2*, *SGTA*, *BCL9L*, *FOXP4*, *KCP*, *MAPK13*, and *PAFAH1B1*) have already been reported to be related to chemotherapy sensitivity (**Fig. 6f**). Specifically, *GPD1* is specifically expressed in brain tumor stem cells and involved in tumor initiation, metastasis, and recurrence; increased expression of *GPD1* correlates with TMZ resistance in GBM^40^. *MSI1* is a stem cell gene that controls the balance between self-renewal and differentiation; GBM lines with *MSI1* knockout display increased sensitivity to cell cycle and DNA replication inhibitors^42^. The rest eight genes, albeit unexplored with GBM chemotherapy, associate closely with gynecological cancers^43–45^, LIHC^46^, gastrointestinal cancers^47, 48^, hematological malignancies^49^, and other cancers^50^, presumably indicating their potential relevance in GBM treatment.

### Cell type identification through GBM feature genes

Evidence has accumulated that GBM exhibits substantial cellular heterogeneity within tumor microenvironment^51^. To examine whether the 167 feature genes identified from bulk multi-omics analysis retain discriminative capacity at single-cell resolution, we further apply MIA to a single-cell dataset (see details in Methods). Based on canonical cell type annotations inferred from established marker genes, the original study classified cells into four cell types^52^ (**Fig. 7a** and **Supplementary Fig. 19a**). Thus, we set the number of clusters at *K*=4 to assess the resolution of MIA clustering, which exhibit a high degree of concordance with the canonical cell type annotations (**Fig. 7b** and **Supplementary Fig. 19b**). Specifically, Clusters 1-4 are predominantly composed of myeloid cells (95.8%), lymphocytes (50.7%), malignant cells (77.9%), and oligodendrocytes (95.0%), respectively (**Supplementary Fig. 19c**). We observe partial co-clustering of lymphocytes and malignant cells in Clusters 2 and 3; when MIA automatically determines the optimal number of clusters as *K*=3, lymphocytes and malignant cells are predominantly divided into the same cluster (**Supplementary Fig. 19d-f** and **Supplementary Table 11**). Trajectory inference reveals a continuous progression spanning myeloid cells, lymphocytes, and malignant cells, with lymphocytes distributed at intermediate pseudo-time values and malignant cells shifted toward later pseudo-time values (**Fig. 7c**), indicating the presence of shared transcriptional programs associated with tumor-immune interactions within the GBM microenvironment^53^.

**Fig. 7.**
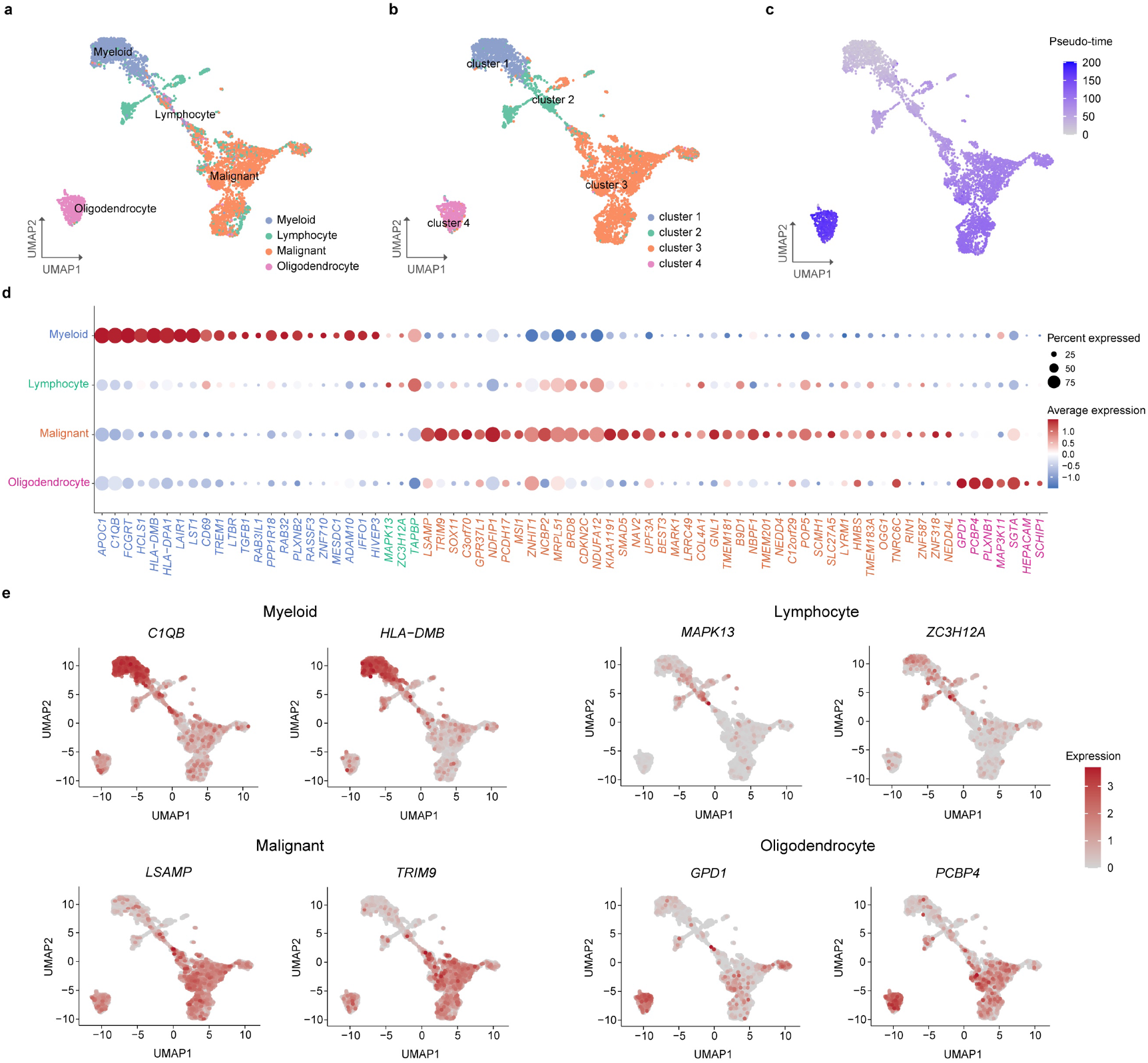
MIA clustering of GBM feature genes in single-cell data. (a) Cell-type annotation based on established marker genes. (b) MIA clustering with the number of clusters fixed at *K*=4. (c) Pseudo-time trajectory of single cells. (d) Differentially expressed feature genes across annotated cell types using a one-versus-rest Wilcoxon rank-sum test (*p*-value<0.01). (e) Top two differentially expressed feature genes in each cell type.

Moreover, among the feature genes, 73 genes exhibit significant differential expression across the four cell types (*p*-value<0.01, Wilcoxon rank-sum test; **Fig. 7d and Supplementary Table 12**). Notably, the highly ranked genes display relatively cell-type-specific expression patterns, with *LSAMP* and *TRIM9* preferentially expressed in malignant cells and *C1QB* and *HLA-DMB* enriched in myeloid cells (**Fig. 7e**). To further assess their biological relevance, we annotate these genes using Cell Taxonomy^54^. Overall, there are 25 genes possessing annotations related to brain tissues, immune cell types, or glioma-related cellular states, in which six genes (*SOX11, MSI1, TREM1, HLA-DMB, C3ORF70* and *HEPACAM*) have been reported to be directly linked to glioma- or GBM-related cell states (**Supplementary Table 12**). Consistent with these annotations, *LSAMP* and *TRIM9* are associated with oligodendrocyte and neural cells, whereas *C1QB* and *HLA-DMB* correspond to microglial/macrophage-associated markers. Notably, the rest 48 genes, lacking cell marker annotations, are found to be enriched in pathways related to receptor tyrosine kinase signaling and oxidative stress responses, indicating that some may represent biologically relevant but currently under-characterized components of GBM cellular heterogeneity (**Supplementary Table 12**). Collectively, these results indicate that MIA is powerful to identify feature genes that capture biologically meaningful cellular heterogeneity at the single-cell level.

## Discussion

In this study, we presented MIA, a tensor-based representation learning framework for multi-omics integrative analysis capable of accurate sample clustering and feature gene identification. Comparative analyses on simulated and real datasets revealed that MIA outperforms existing algorithms in both integrative clustering and feature discovery. As testified in GBM multi-omics data analysis, MIA is powerful to cluster samples into clinically relevant subtypes with distinct survival outcomes and identify feature genes associated with TMZ treatment sensitivity. MIA can also be applied to single-cell data, where the identified feature genes retain discriminative capacity across major GBM cell populations at single-cell resolution.

MIA adopts a machine learning-based strategy to achieve powerful multi-omics integrative analysis. Although deep learning has achieved remarkable progress in biomedical research, its reliance on large, well-annotated datasets limits its applicability to omics data, which are typically high-dimensional, unlabeled, and sample-limited. In contrast, the machine learning models used in MIA, such as random forest, are more robust to small sample size, less prone to overfitting, and offer interpretable results with biological insights. Despite this, technically, future explorations and improvements for MIA include the following. First, the tensor framework requires matched samples and corresponding features across omics types, limiting its applicability to data with heterogeneous features. Second, after Tucker decomposition, only sample-feature matrix is used for clustering, whereas gene-feature and omics type-feature matrices remain underutilized. Notably, clustering heatmaps from gene-feature matrix in simulated data clear separated real from noise features (**Supplementary Fig. 1b**), suggesting great potential for feature filtering. Additionally, omics type-feature matrix may quantify contributions from different omics layers; in GBM multi-omics, for example, eigenvalues for CNV, gene expression, and methylation were 0.446, 0.674, and 0.589, respectively (**Supplementary Table 7**), indicating a relatively larger contribution of gene expression to the learned representation, which aligns well with single-omics analyses showing its most relevance to survival (**Fig. 3c**). Thus, omics type-feature matrix contains substantial biological relevance and further exploration of its application for optimizing feature gene identification would be desirable.

Across the ten cancer types, clusters derived from CNV and methylation data generally showed smaller survival differences than those from gene expression (**Fig. 3c**). From the data perspective, this may stem from limited or/and uneven distribution of samples exhibiting heterogeneous copy number amplifications/deletions, and signal attenuation introduced during the conversion of methylation measurements from the site level to the gene level. From the biological perspective, gene expression is closely correlated with CNV and promoter methylation status^55^, together exerting an influence on protein production, cell function, and thereby, phenotypic and clinical outcomes^56^. Furthermore, we observe that nearly all algorithms exhibit comparable survival differences in LGG on multiple omics combinations, primarily due to substantial molecular variations between grade II and grade III LGG, with the latter associated with a worse prognosis^57^.

In GBM, we observed that the fraction of genome altered, a measure of the extent of copy number changes in the genome, decreased with declining survival rates (**Fig. 4d**). In early-stage tumors, studies have demonstrated that a higher fraction of genome altered is typically associated with poorer pathological characteristics and survival rates^58, 59^, and is also a sign of poor prognosis^60, 61^. On the contrary, in advanced tumors, several studies have also found that tumors with lower genomic alterations can exhibit more aggressive behavior and decreased survival rates^33^. For example, in metastases compared to primary lung adenocarcinoma, the fraction of genome altered was significantly lower^62^. Additionally, excessive genomic instability may cumulate damage, exceeding the threshold of tumor cell activity, thereby inhibiting advanced tumor growth and metastasis^63^. Our observations in GBM are consistent with these findings of advanced tumors, but more research is needed to explore the association between this indicator and the prognosis of advanced tumors. Furthermore, single-cell analysis demonstrates that the feature genes identified by MIA remain informative at cellular resolution, supporting their biological relevance across both bulk and single-cell data. However, MIA shows reduced resolution in separating closely related cell populations, as evidenced by the partial overlap between lymphocyte and malignant cells. Consequently, MIA may currently be better suited for feature discovery than for fine-grained cell-type annotation, while future adaptation to single-cell multi-omics analyses may further enhance its ability to characterize cellular heterogeneity.

To sum up, with the vast volume of multi-omics data generated by higher-throughput sequencing technologies, multi-omics integrative analysis becomes increasingly routine and popular. Thus, MIA bears great utility for providing multi-layered insights into complex biological and physiopathological processes and promoting precision diagnostics and targeted therapies.

## Methods

### Simulated datasets

Simulated multi-omics datasets were generated using a custom Python script. Five parameters—number of clusters (*K*), number of omics types (*T*), sample size (*N*), number of real features (*M_t_*), and number of noise features (*M_f_*)—were considered and five replicates were generated for each parameter combination. Samples were assigned to *K* clusters using a Gaussian probability distribution. Each omics layer was represented as an *N*×(*M_t_*+*M_f_*) matrix, where *M_t_* real features were differentially expressed across clusters with group-specific means and Gaussian noise, and whereas *M_f_* noise features were randomly generated. For clustering evaluation, the effects of *K* and *T* were first evaluated (*K*=2∼6 and *T*=1∼3 with *N*=500, *M_t_*=5,000 and *M_f_*=5,000 fixed), and followed by evaluation of the effects of *N*, *M_t_*, and *M_f_* (*N*=50, 500, or 5,000, *M_t_*=1,000, 5,000, or 10,000, and *M_f_*=1,000, 5,000, or 10,000, with *K*=4 and *T*=2 fixed). For feature gene identification, datasets were simulated with *K*=4 or 5, *N*=500, *M_t_*=1,000, and *M_f_*=10,000. The simulation script was provided as *simulate_datasets.py*in the MIA package, available at https://ngdc.cncb.ac.cn/biocode/tool/BT008143.

### Empirical datasets from TCGA

Multi-omics datasets from ten TCGA cancer types, including CNV (gene-level), gene expression (FPKM), and DNA methylation, were downloaded from UCSC Xena (https://xenabrowser.net/). For DNA methylation data, gene-level methylation values were calculated by averaging the methylation levels of all probes located within 1,500 bp upstream of each gene. Only samples and genes shared across all three omics types were retained for integrative analysis. Survival data and clinical information were obtained from cBioPortal (https://www.cbioportal.org/). The mRNA expression-based stemness index (mRNAsi) was obtained from a previous study ^34^. Feature genes identified for GBM were annotated by PubMed (https://pubmed.ncbi.nlm.nih.gov/), BrainBase^64^, MSigDB^65^, and EWAS Atlas^66^.

### GBM temozolomide treatment datasets

Gene expression datasets related to GBM response to TMZ treatment were retrieved from the Gene Expression Omnibus (GEO) (https://www.ncbi.nlm.nih.gov/gds/). Specifically, GSE211272^41^ contained 72 samples derived from 12 GBM cell lines, including 24 control samples, 24 samples treated with TMZ and radiotherapy, and 24 samples treated with TMZ, radiotherapy and molecular inhibitor of Ataxia-telangiectasia and Rad3-Related protein. GSE113510^67^ comprised three control samples and three TMZ-resistant GBM cell samples. GSE100736 included three TMZ-resistant GBM cell lines and 3 TMZ-sensitive cell lines. GSE68029^68^ consisted of three samples surviving one cycle and three samples surviving two cycles of treatment with 500µM TMZ, representing low to high TMZ resistance. Batch effects were corrected using Combat^69^ method. Expression matrices of the predefined 167 GBM feature genes were then generated by mapping these genes to each dataset, resulting in 165, 161, 165, and 162 matched genes in GSE211272, GSE113510, GSE100736, and GSE68029, respectively.

### Single-cell data processing and analysis

Single-cell RNA-sequencing data were obtained from the Chinese Glioma Genome Atlas (CGGA), including 11 GBM samples, 5,124 cells, and 24,989 genes^70–72^. Batch effects were corrected using Seurat^73^ with SCTransform normalization and RPCA-based integration, after which 119 of the 167 predefined feature genes were mapped to the dataset and used for MIA clustering. Differentially expressed genes associated with cell types were identified using the FindAllMarkers function in Seurat. Significant genes met the following criteria: (1) expression in ≥20% of cells in at least one cluster; (2) log_2_ fold change≥0.5; and (3) adjusted Wilcoxon rank-sum test *p*-value<0.01. Following dimensionality reduction with Harmony^74^, the embeddings were visualized using UMAP, and cellular trajectories were reconstructed using Slingshot^75^.

### Tensor integration and decomposition

Multi-omics data were represented as a three-dimensional tensor *A* ∈ *R^N×M×T^*, where *N* is the number of samples, *M* is the number of genes, and *T* is the number of omics types. For unlabeled data, Tucker decomposition was applied to decompose *A* into a core tensor and three feature matrices:

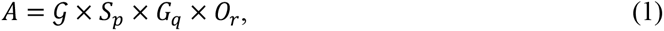

where *S* ∈ *R^N×P^*, *G* ∈ *R^M×q^* and *O* ∈ *R^T×r^* denote the sample-, gene-, and omics-feature matrices, respectively, and G ∈ *R^p×q×r^* is the core tensor describing the interactions among the latent components. The dimensions *p*, *q*, and *r* satisfy *p*≤*N*, *q*≤*M*, and *r*≤*T* and are set to *log_2_*(*N*), *log_2_*(*M*) and *log_2_*(*T*) by default, respectively, and they can also be configured to specific values by users. The sample-feature matrix *S* was subsequently used for clustering analysis. Tucker decomposition was implemented using TensorLy^76^.

### FCM clustering and parameter optimization

The sample-feature matrix *S* obtained from Tucker decomposition was used for FCM clustering, partitioning *N* samples into *K* (*K*>1) fuzzy clusters by minimizing the following objective function:

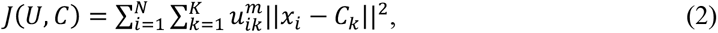

where *u_ik_* ∈ [0,1] (*i*=1, 2, …, *N* and *k=*1, 2, …, *K*) is the membership matrix, *C_k_* (*k=1*, 2, …, *K*) denotes the cluster centers, and *m* ∈ (1, ∞) is the fuzzy index. The *u_ik_* and *C_k_* are updated iteratively according to the standard FCM algorithm until convergence, defined as *J_n_-J_n-1_<e^-^*^15^.

Two parameters were optimized for FCM: the fuzzy index (*m*) and the number of clusters (*K*). The optimal *m* was estimated using Schwammle’s formula^29^: *m=*1+(1418/*N*+22.05)*M*^(−2)+(12.33/*N*+0.243)*M*^(−0.0406*ln*(*N*)−0.1134), where *N* is the number of samples and *M* is the number of features. When *K* was not specified, it was automatically determined using five clustering evaluation metrics, including the Silhouette Coefficient^77^, Davies-Bouldin Index^78^, Elbow method^79^, Separation index, and Compactness index. Among these metrics, the Silhouette Coefficient consistently achieved the highest accuracy for estimating the optimal number of clusters (**Supplementary Fig. 3**) and was therefore used to determine the optimal *K*.

### Gene filtering by Gini index

The Gini index was used to filter genes prior to ERFM analysis (**Supplementary Fig. 4a,b**). For the *t*-th omics type, the Gini index of gene *j* under a partition threshold *X* was calculated as:

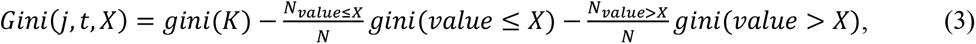

where *N* is the total number of samples, and *value*≤*X* (or *value*>*X*) denotes the numbers of samples with values less than or equal to (or greater than) threshold *X*. The objective is to identify the optimal threshold *X* that maximizes *Gini*(*j,t,X*):*Gini*(*j,t*)=*max*(*Gini*(*j*,*t,X_1_*), *Gini*(*j,t,X_2_*), …, *Gini*(*j,t,X_n_*)). The mean Gini index for the *t*-th omics type was then calculated as

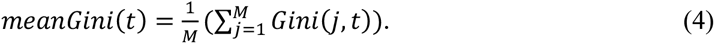

Genes satisfying the condition *Gini*(*j,t*)>*meanGini*(*t*) across all omics types were retained. These retained genes, denoted as *M’*, were considered as potential features for subsequent ERFM analysis.

### Feature gene identification based on multi-omics using ERFM

In this study, ERFM, an enhanced RFM, was developed to identify feature genes. After gene filtering, ERFM was constructed using the retained genes, multi-omics data, and the corresponding sample labels or clustering results (**Supplementary Fig. 4c**):

**(a) Data preparation**: Using bootstrap sampling, 70% of samples are randomly selected for training each decision tree, while the remaining 30% of samples are used for testing. Each decision tree has different training and testing samples. *M_s_* (*M_s_*≤*sqrt*(*M’*)) genes are randomly selected from the *M’* genes, and the Gini index is used to select the best combination among the *M_s_* genes for constructing the decision tree.

**(b) Enhanced tree structure**: Unlike conventional decision trees, in which each node represents a single feature, ERFM simultaneously incorporates all omics layers of an individual gene into the tree structure. Specifically, once a gene *M_i_* was selected, data from all *T* omics layers of that gene are used to construct tree nodes, yielding *T* nodes per gene. The optimal splits for each node are determined using *Gini*(*j*,1), *Gini*(*j*,2), …, and *Gini*(*j*, *T*). For example, in the case of three omics types, three nodes are built for a single gene. The trees are grown until either the depth reaches *log*2(*M’*)-1 or the number of features reaches *sqrt*(*M’*).

**(c) Gene importance in a decision tree**: Once a decision tree *d* is trained, the remaining 30% of samples are used for classification prediction to calculate out-of-bag error rate (*obb_error_d_*). To evaluate the importance of a gene *M_i_*, the values of all omics layers corresponding to *M_i_* in the test set are simultaneously randomized, and the out-of-bag error rate (*obb_error*_random(*Mi*)_) is calculated. The importance of gene *M_i_* in the decision tree *d* is then defined as: *importance_d_*(*M_i_,d*)=*obb_error*_random(*Mi*)_–*obb_error_d_*.

**(d) Gene importance in a random forest**: A random forest *r* creates 100 decision trees *d*∈{1, 2, …,100} by default, repeating **steps (a-c)**. The importance of gene *M_i_* in forest *r* is calculated as: 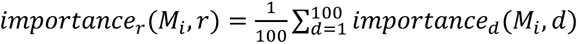. The number of genes whose *importance_r_*(*M_i_,r*)≤0 is recorded as *M_≤_*_0_.

**(e) Iterative computational procedure**: To enhance consistency of the identified feature gene set, an iterative procedure is applied with a termination condition: if *M_≤_*_0_*>0*, these *M_≤_*_0_ genes are removed, and *M’=M’-M_≤_*_0_ is updated to repeat **steps (a-d)**; if *M_≤_*_0_*=0*, the iterative procedure stops, and the remaining *M’* genes are considered as the final feature gene set. The overall importance of gene *M_i_* is computed as: 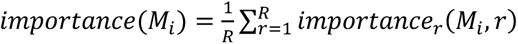, where *R* represents the number of iterations.

Finally, MIA normalizes and ranks the importance scores of the remaining *M’* genes, and the Wilcoxon rank-sum test is used to identify genes showing significant differences among clusters (**Supplementary Fig. 4d**).

### Clustering evaluation

To benchmark the performance of MIA, all algorithms were applied to the same preprocessed multi-omics datasets, and clustering results were evaluated using a unified evaluation framework to ensure consistency and comparability. For simulated datasets, clustering accuracy was assessed using the ARI, which measures the agreement between predicted and true cluster assignments. For TCGA empirical datasets, the biological relevance of the identified clusters was evaluated using survival and clinical analyses. Overall survival was assessed using Kaplan-Meier analysis with log-rank tests. Associations between clusters and categorical clinical records were evaluated using the chi-square test, and continuous variables were analyzed using the Kruskal-Wallis test. Statistical significance was further assessed by permutation testing to estimate empirical confidence intervals, following previously established procedures^12^. For TMZ response datasets, clustering performance was evaluated using multi-class ROC analysis in a one-vs-rest framework. Overall classification performance was summarized by the macro-averaged area under the ROC curve (AUC).

### Feature gene evaluation

The significance of individual feature genes was evaluated using the Kruskal-Wallis test. For each gene, significance was calculated independently in each omics layer using known sample labels in both simulated and empirical datasets. The resulting significance values were then averaged across the corresponding omics layers to obtain a multi-omics significance score for each gene. To evaluate the clustering power of an identified feature gene set, the multi-omics significance scores and Gini indices were averaged across all feature genes, yielding overall clustering scores for different feature gene sets.

## Data availability

All datasets used in this study are publicly available. Details are provided in the **Methods** section.

## Code availability

MIA v1.0 was used to generate the results in this work and is freely available for academic research use at https://ngdc.cncb.ac.cn/biocode/tool/BT008143, and the web server can be accessed at https://ngdc.cncb.ac.cn/mia/.

## Acknowledgments

This work was supported by the National Natural Science Foundation of China (T2425005 and 82541014 to ZZ), Youth Innovation Promotion Association of the Chinese Academy of Sciences (Y2023027 to LM), CNCB-initiative program (CNCBi2025001 to LM and ZZ).

## Author contributions

Yang Zhang: Data curation, Formal analysis, Investigation, Methodology, Software, Visualization, Validation, Writing - original draft, Writing - review & editing; Lin Liu: Data curation; Zhaoqi Liu: Writing - review & editing; Qi Liu: Writing - review & editing; Lina Ma: Conceptualization, Project administration, Funding acquisition, Writing - review & editing; Zhang Zhang: Conceptualization, Funding acquisition, Project administration, Writing - review & editing. All authors have read and approved the final manuscript.

## Supplementary Figures

**Supplementary Fig. 1.**
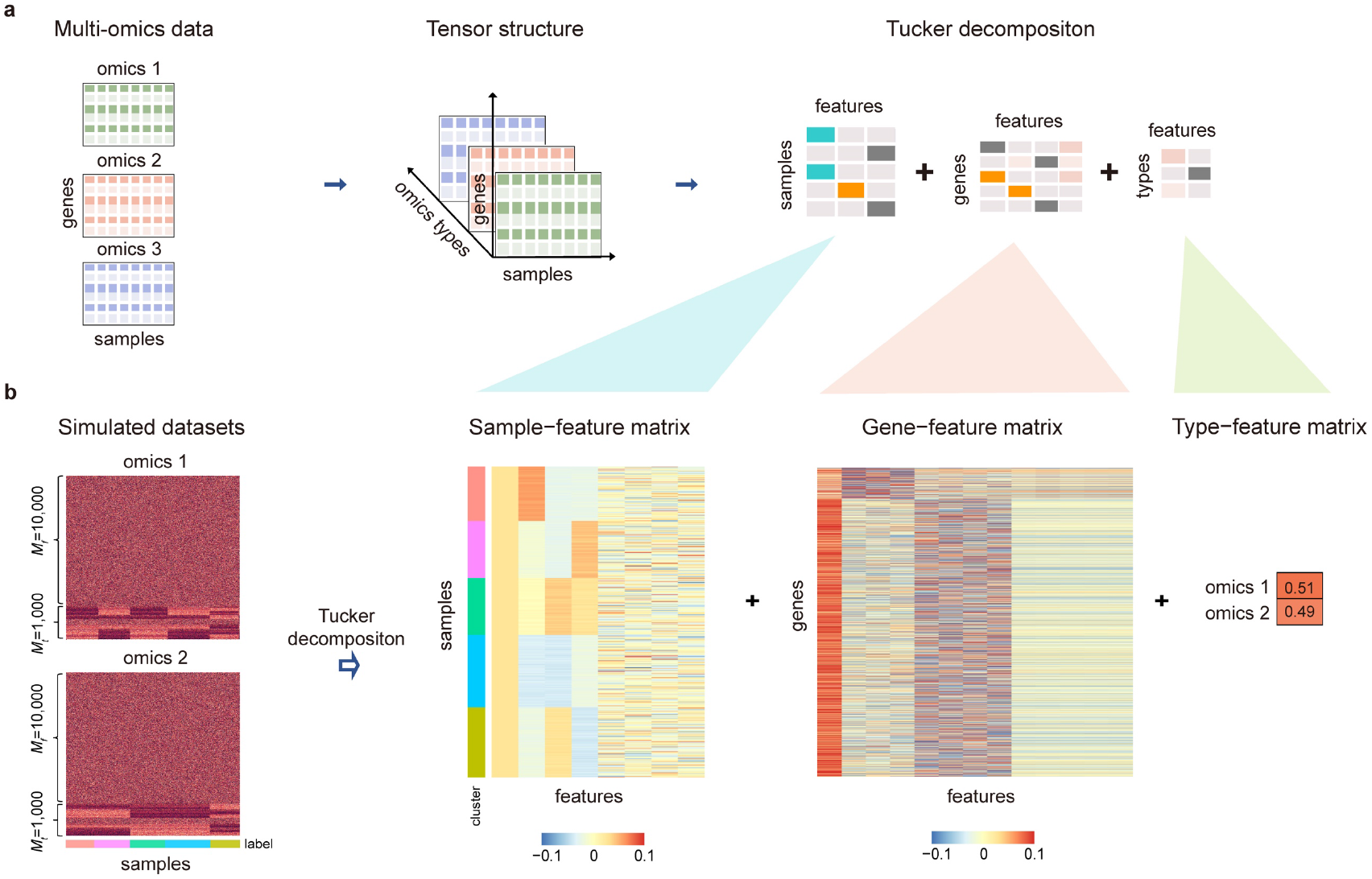
Schematic of Tucker decomposition for multi-omics data. (a) Tensor integration of multi-omics data into a three-dimensional tensor, and Tucker decomposition factorizing the tensor into three factor matrices: sample-feature matrix, gene-feature matrix, and type-feature matrix. (b) Tucker decomposition results on simulated datasets with *T*=2, *K*=5, *N*=500, *Mt*=1,000, and *Mf*=10,000.

**Supplementary Fig. 2.**
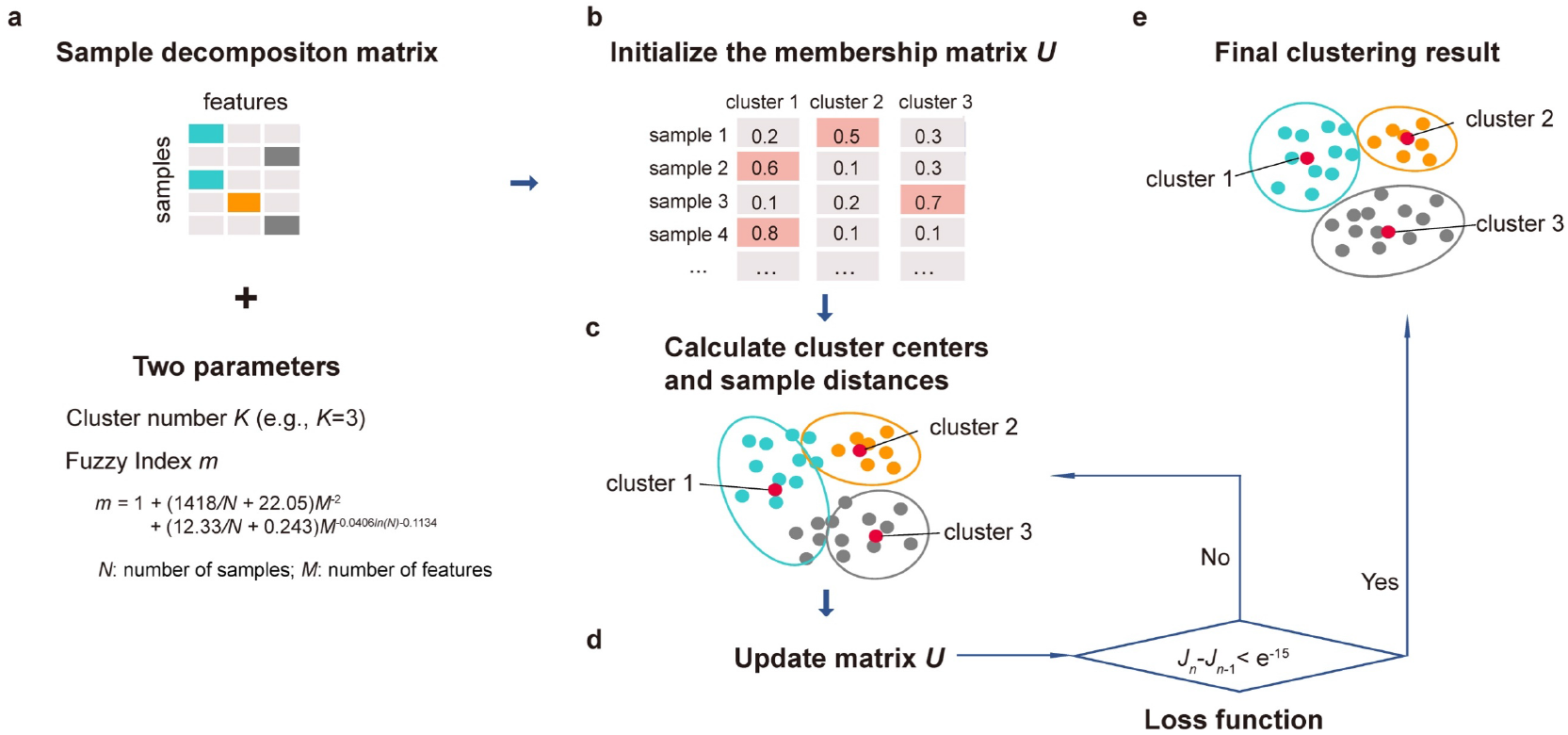
Schematic of multi-omics data integration and FCM clustering. (a) FCM clustering using the sample-feature matrix with specified cluster count (*K*) and fuzzy index (*m*). (b) Membership matrix *U* initialized to represent sample-cluster membership probabilities. (c) Cluster center vectors calculated from *U*. (d) *U* iteratively updated based on cluster centers and *m*, stopping when the objective function converges below a threshold. (e) Each sample assigned to the cluster with the highest membership probability in *U*.

**Supplementary Fig. 3.**
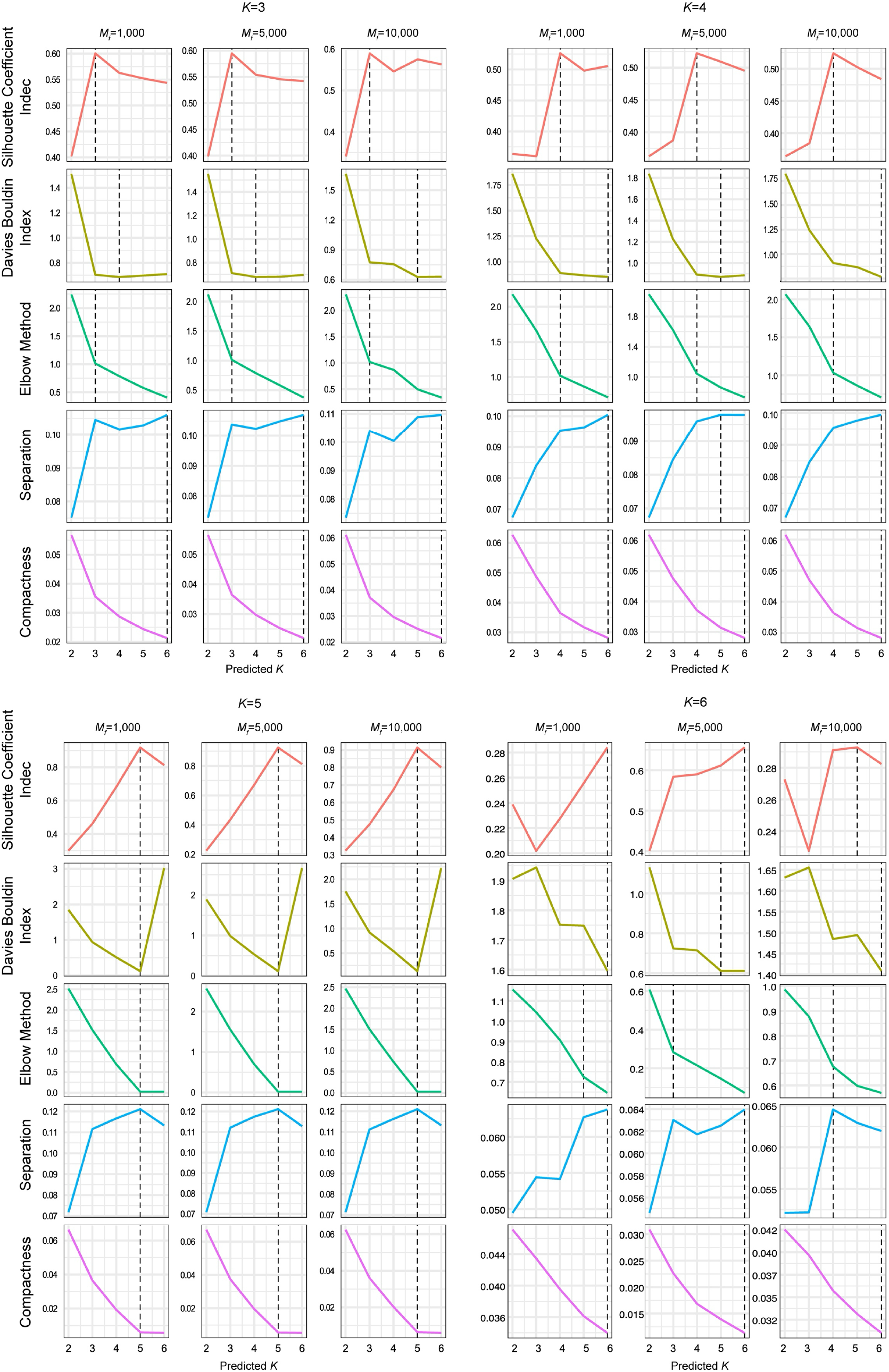
Evaluation of five clustering evaluation metrics for optimal *K* selection on simulated datasets when *K* varies from 3 to 6. Simulated datasets comprise *K*=3, 4, 5, 6 and *Mf*=1,000, 5,000, and 10,000, with fixed *N*=500 and *Mt*=1,000. The optimal *K* values are evaluated by five indicators: Silhouette Coefficient, Davis Bouldin Index, Elbow method, Separation index, and Compactness index.

**Supplementary Fig. 4.**
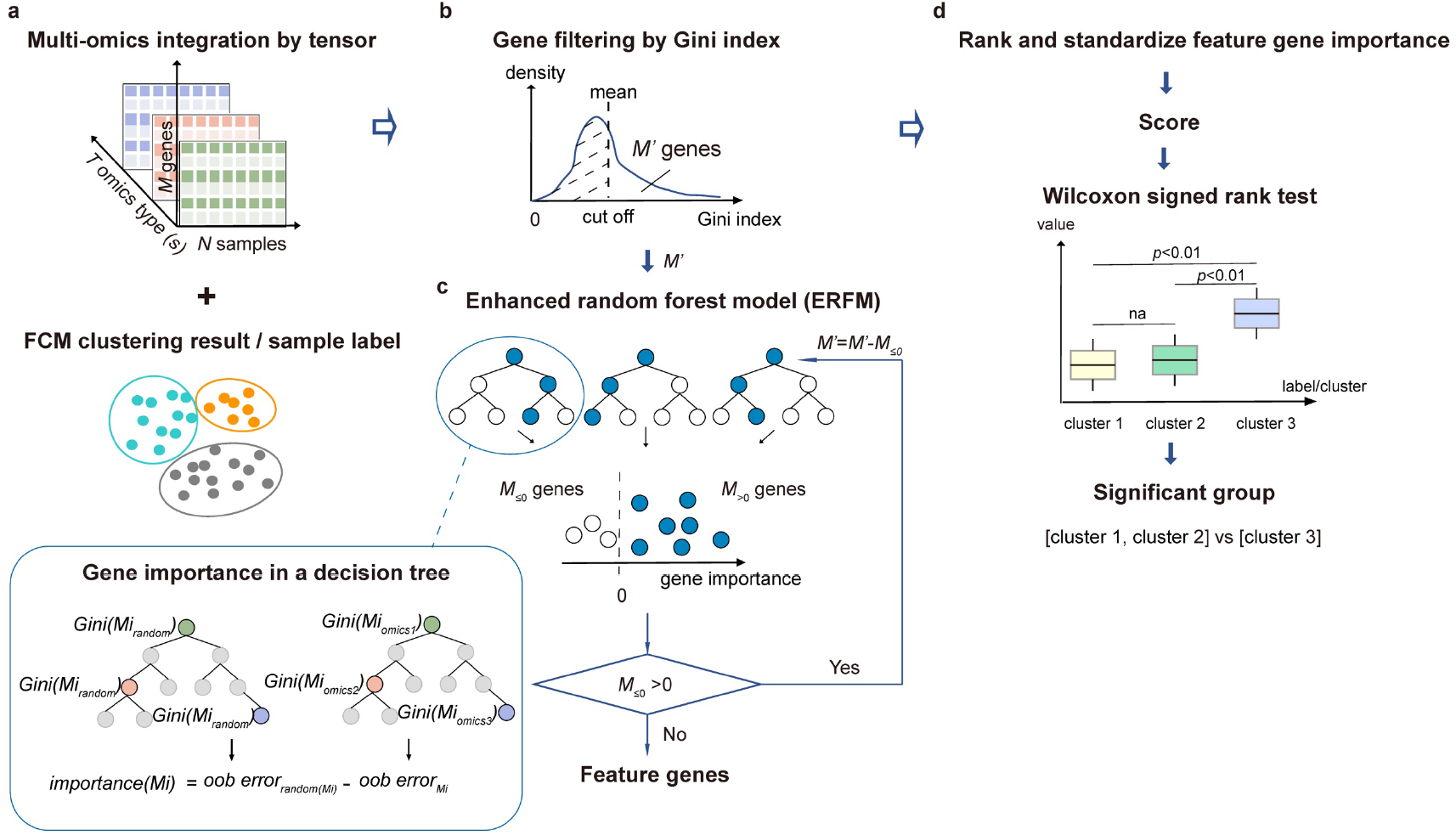
Feature gene identification by ERFM. (a) Input multi-omics data with corresponding clustering results or sample labels. (b) Preliminary gene filtering based on the Gini index. Genes with values exceeding the mean across all omics are retained. (c) Construction of the ERFM adapted for three-dimensional tensors. Gene importance is quantified by the reduction in classification accuracy following simultaneous randomization of all omics types for each gene. The computational procedure iterates until all genes have positive importance values. (d) Final feature-gene scoring, in which importance values are normalized and significant differences across clusters are evaluated using the Wilcoxon rank-sum test (see details in Methods).

**Supplementary Fig. 5.**
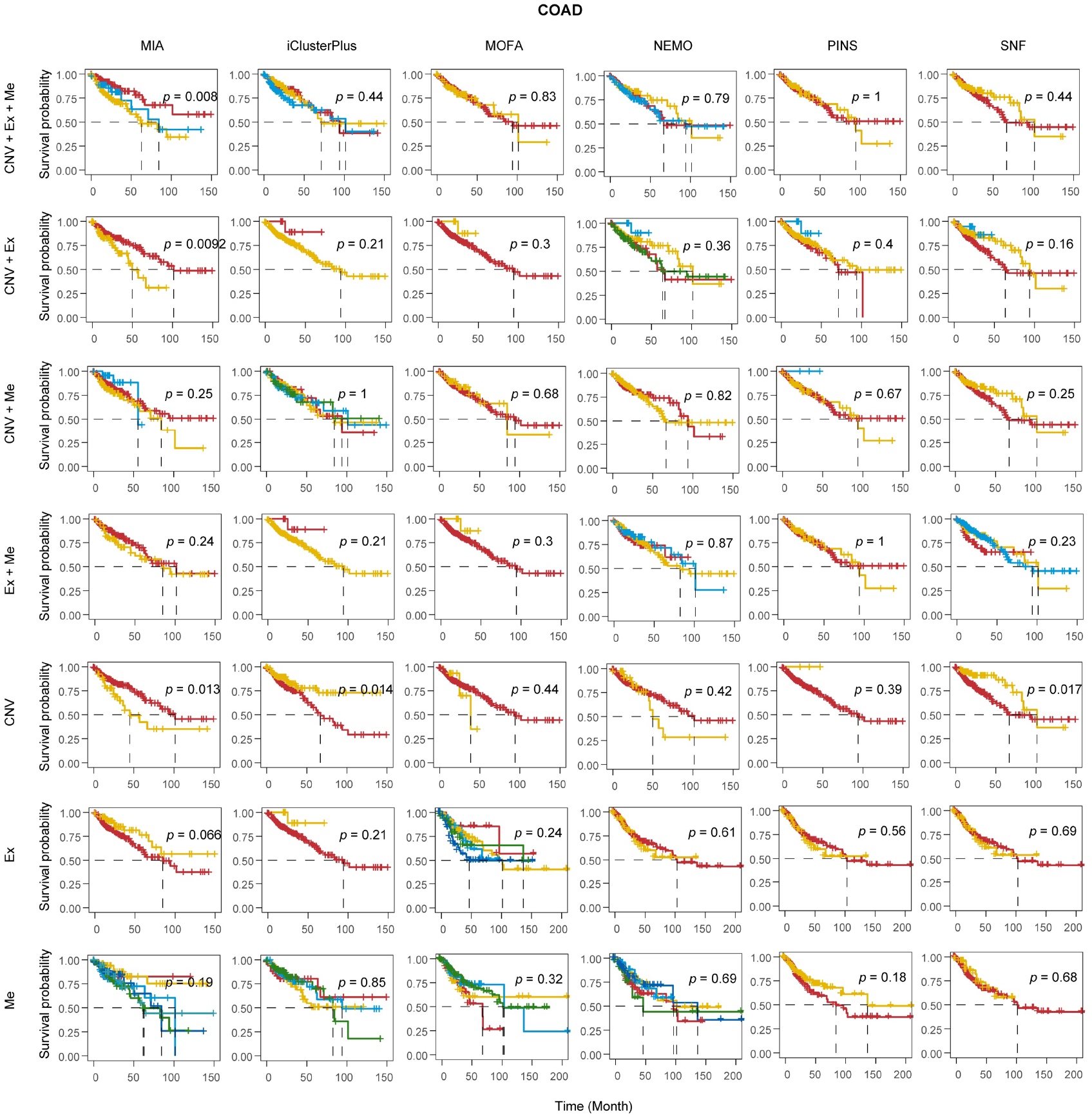
Kaplan-Meier survival analysis on TCGA COAD dataset clustered by different algorithms across various omics combinations. The log-rank test was used to assess the statistical significance between survival curves. Abbreviations: CNV, copy number variation; Ex, gene expression; Me, DNA methylation.

**Supplementary Fig. 6.**
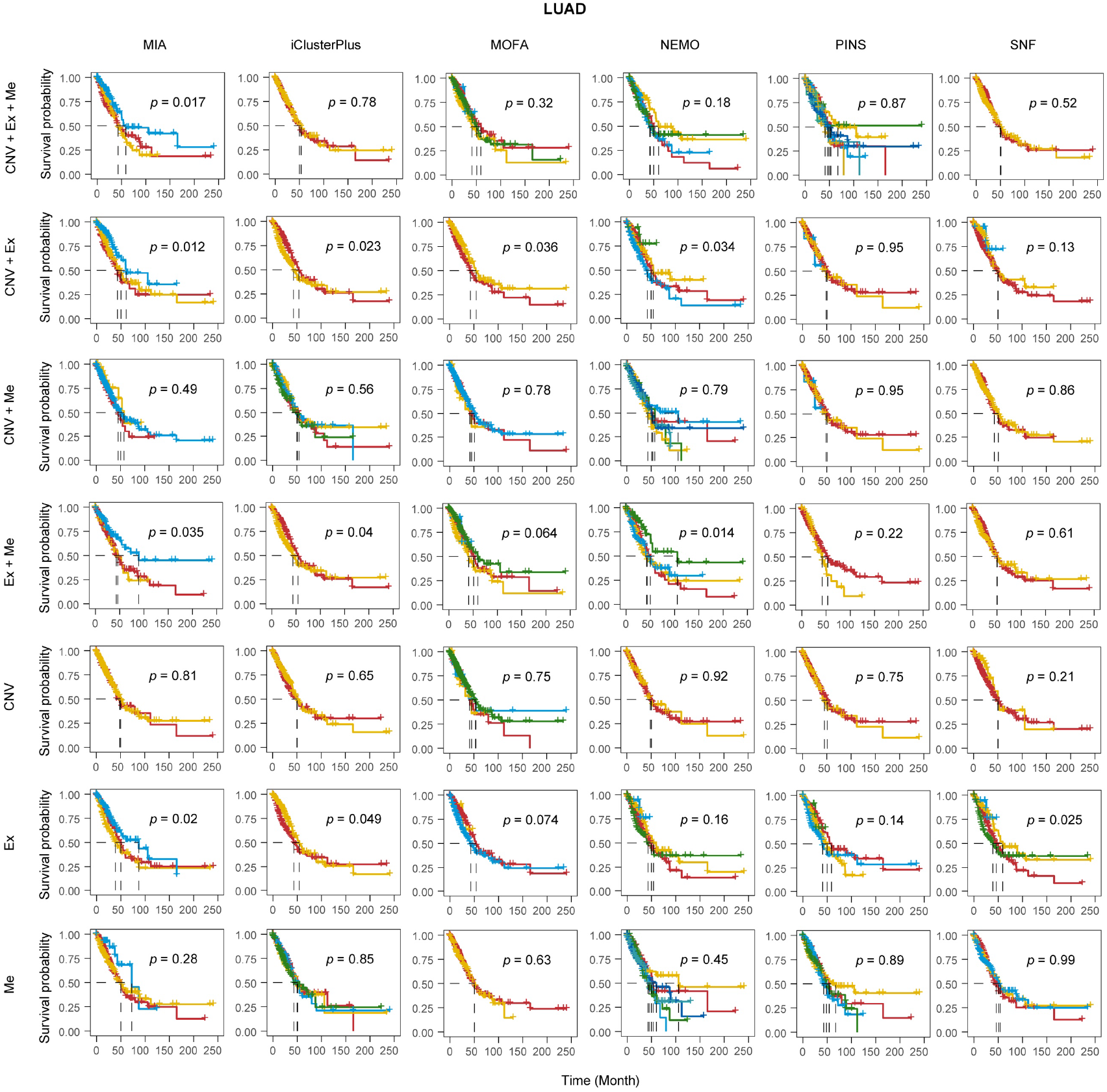
Kaplan-Meier survival analysis on TCGA LUAD dataset clustered by different algorithms across various omics combinations. The log-rank test was used to assess statistical significance between survival curves. Abbreviations: CNV, copy number variation; Ex, gene expression; Me, DNA methylation.

**Supplementary Fig. 7.**
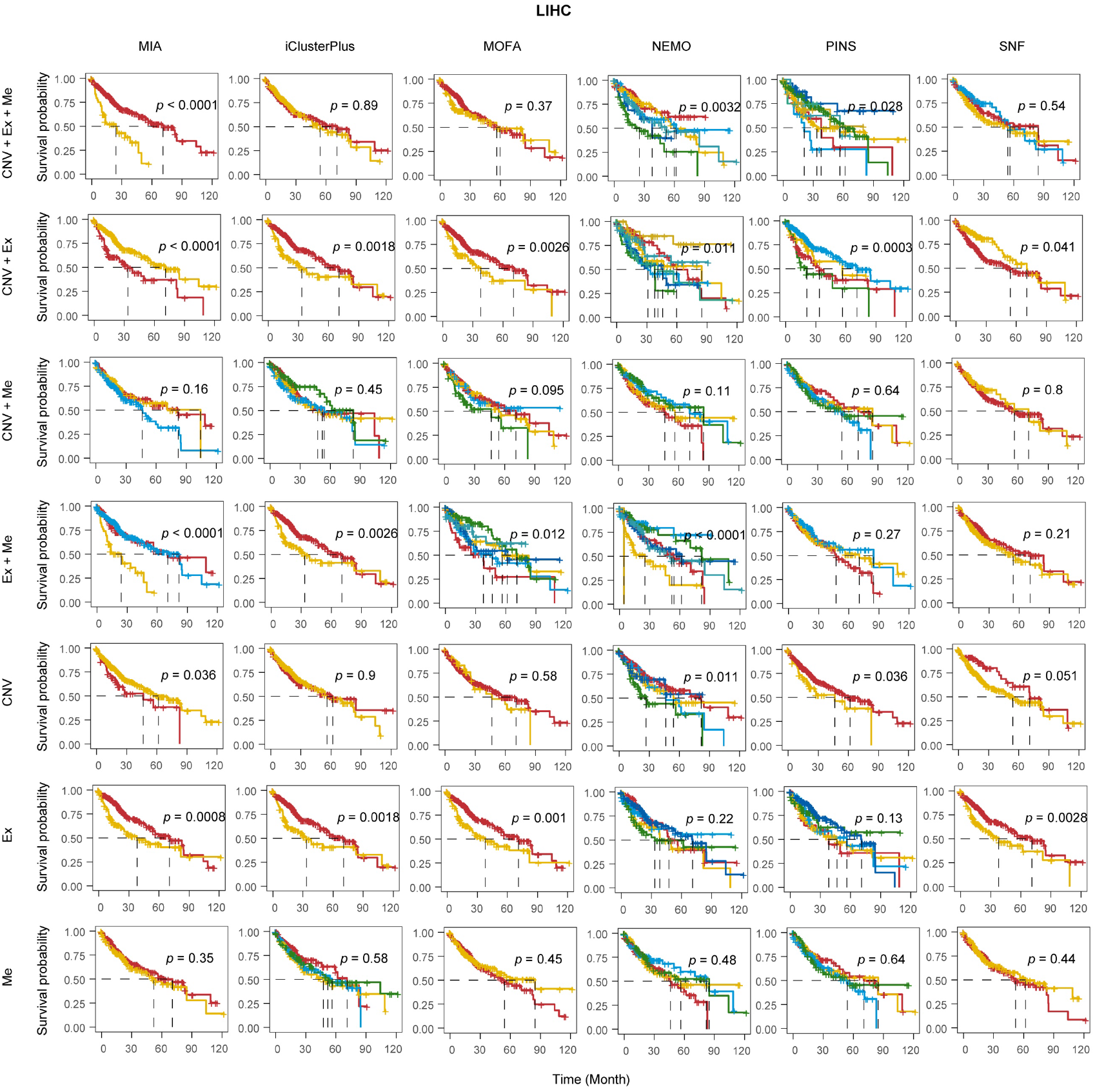
Kaplan-Meier survival analysis on TCGA LIHC dataset clustered by different algorithms across various omics combinations. The log-rank test was used to assess statistical significance between survival curves. Abbreviations: CNV, copy number variation; Ex, gene expression; Me, DNA methylation.

**Supplementary Fig. 8.**
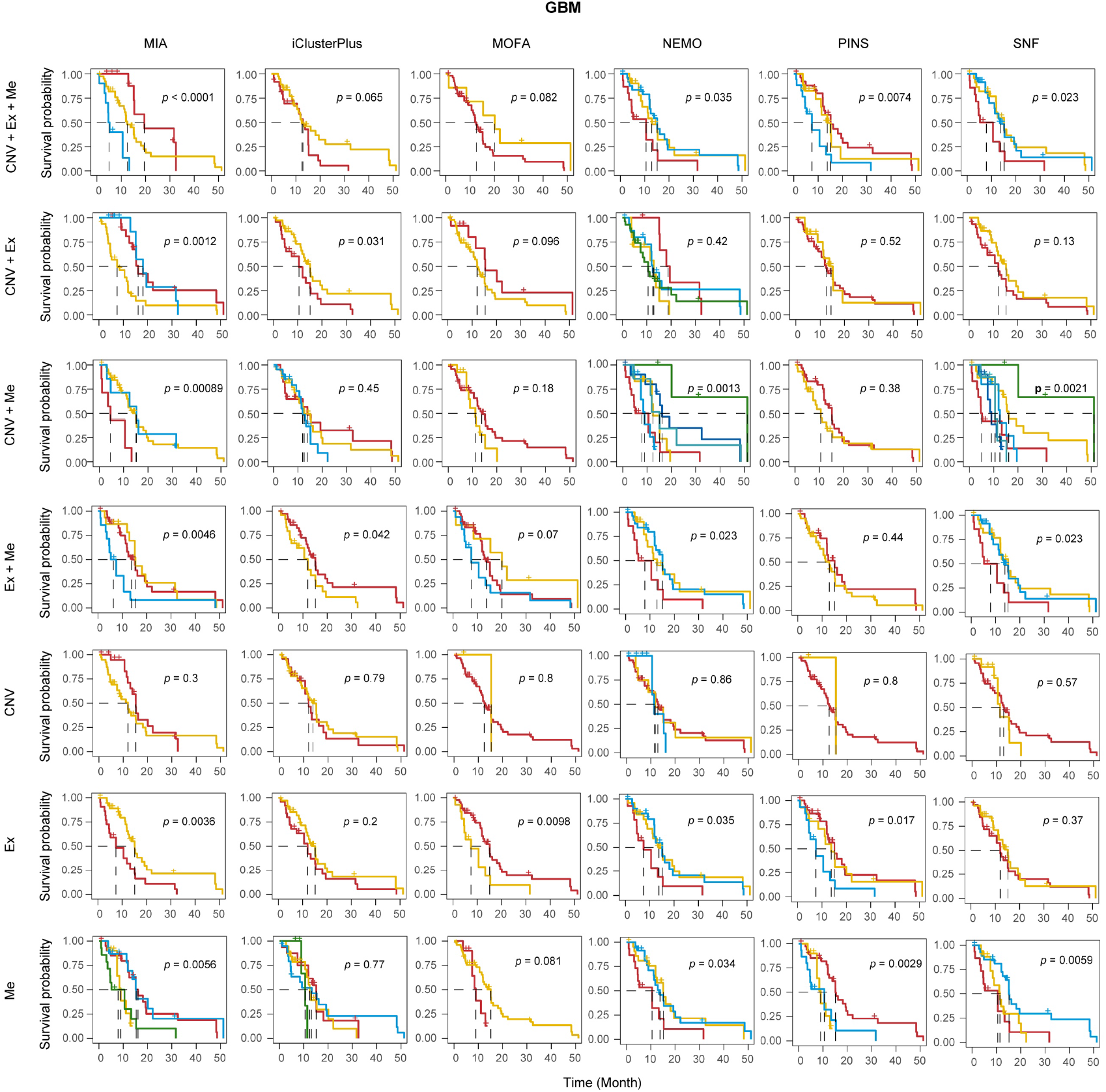
Kaplan-Meier survival analysis on TCGA GBM dataset clustered by different algorithms across various omics combinations. The log-rank test was used to assess statistical significance between survival curves. Abbreviations: CNV, copy number variation; Ex, gene expression; Me, DNA methylation.

**Supplementary Fig. 9.**
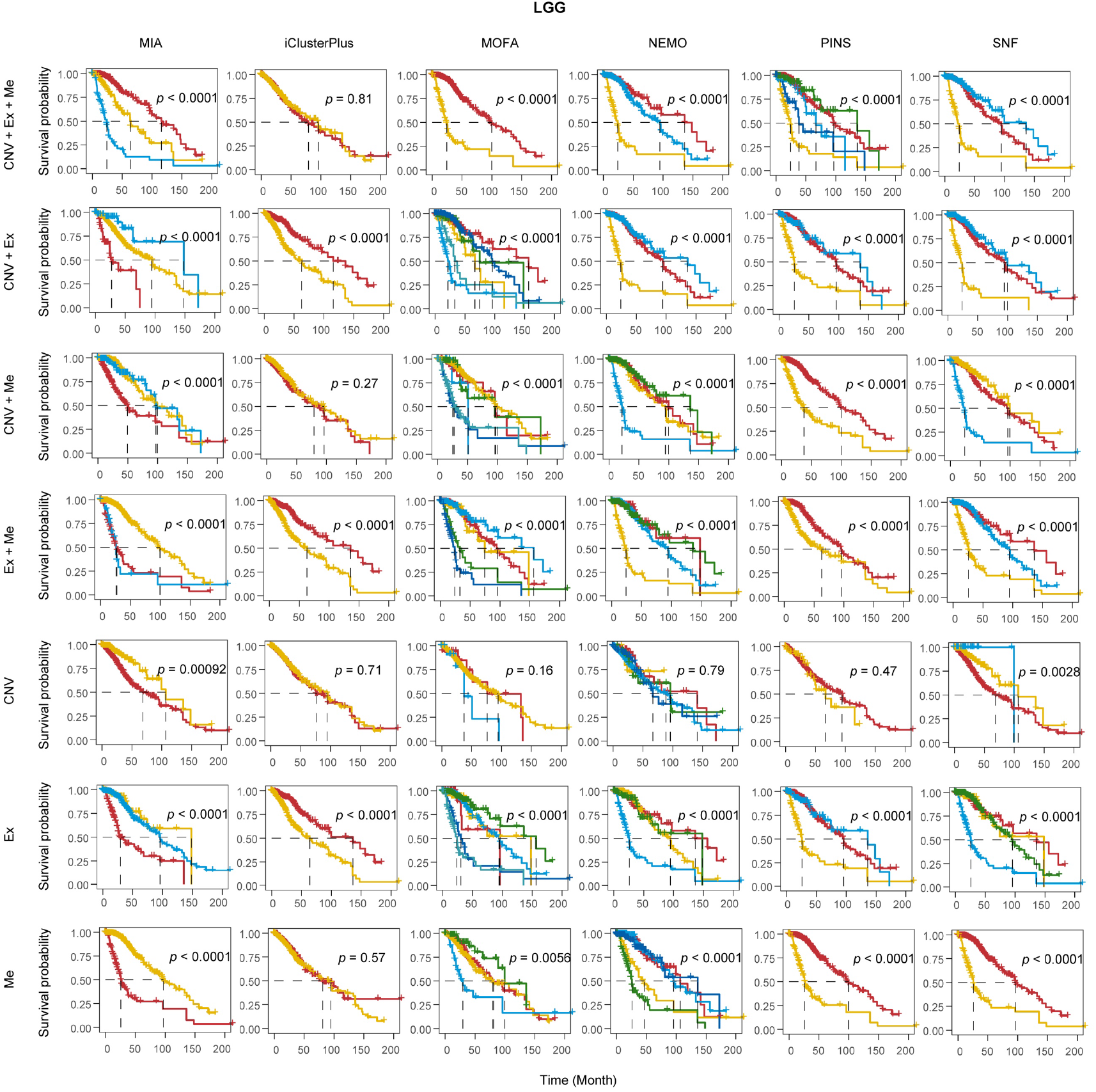
Kaplan-Meier survival analysis on TCGA LGG dataset clustered by different algorithms across various omics combinations. The log-rank test was used to assess statistical significance between survival curves. Abbreviations: CNV, copy number variation; Ex, gene expression; Me, DNA methylation.

**Supplementary Fig. 10.**
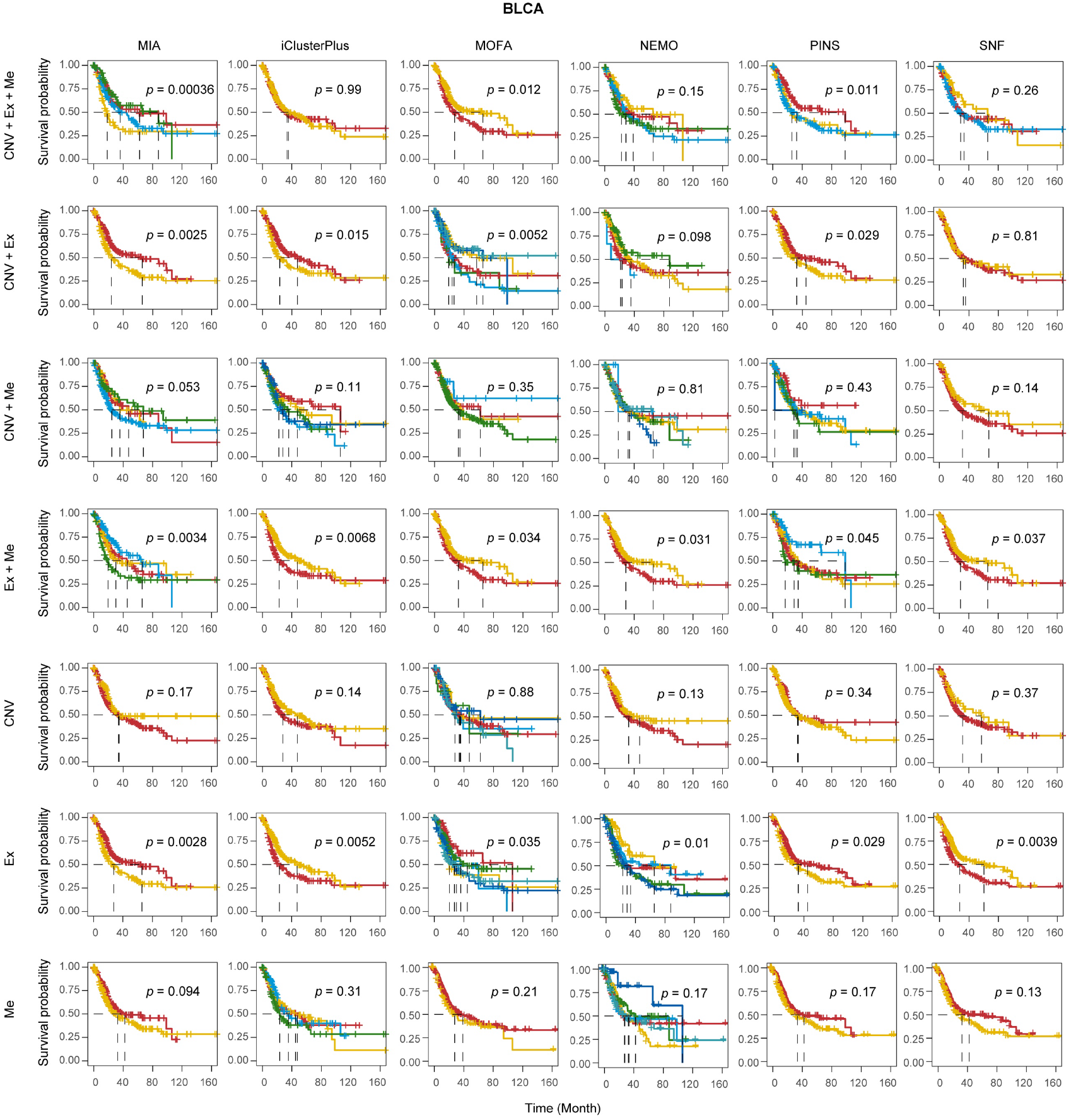
Kaplan-Meier survival analysis on TCGA BLCA dataset clustered by different algorithms across various omics combinations. The log-rank test was used to assess statistical significance between survival curves. Abbreviations: CNV, copy number variation; Ex, gene expression; Me, DNA methylation.

**Supplementary Fig. 11.**
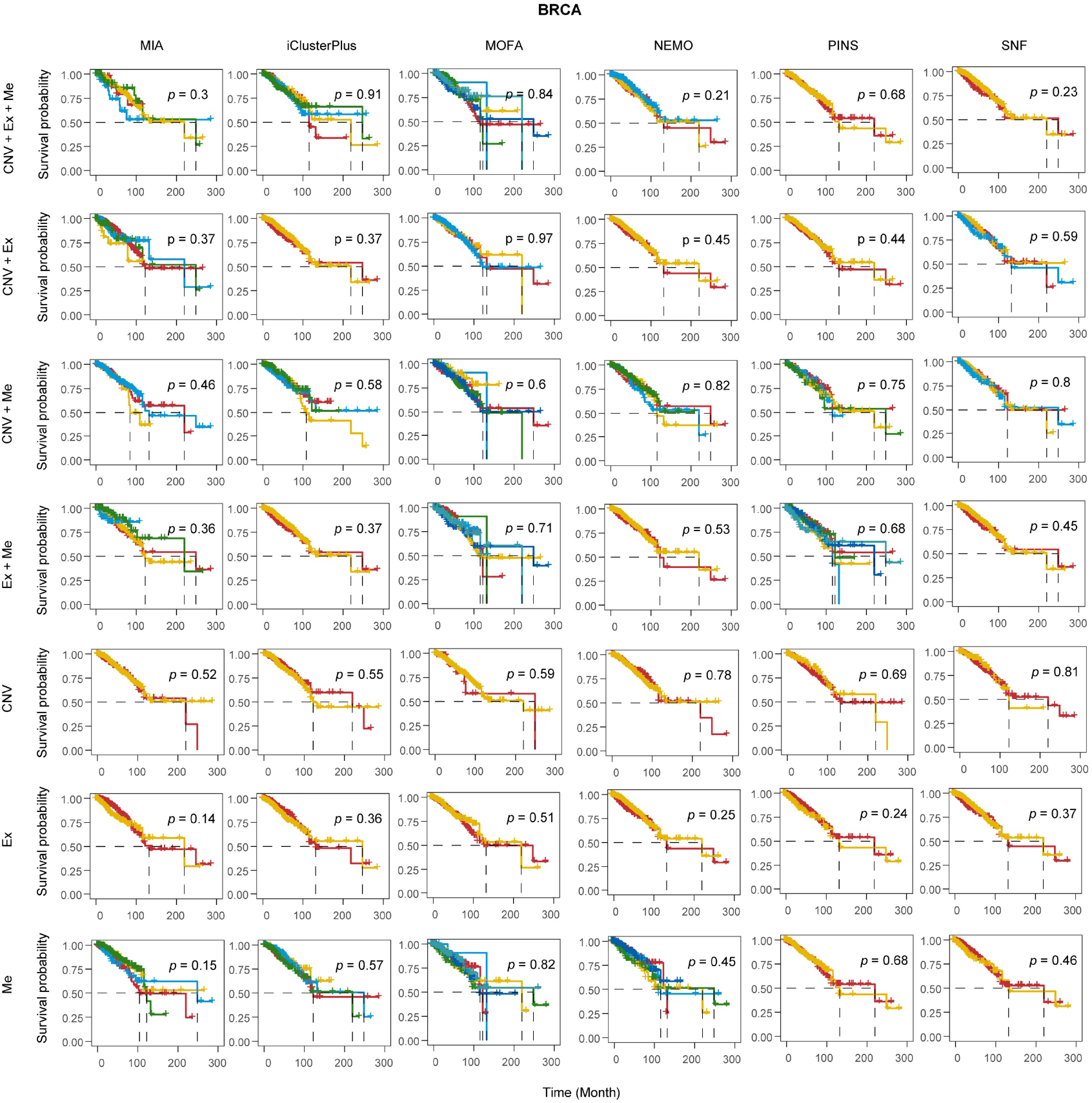
Kaplan-Meier survival analysis on TCGA BRCA dataset clustered by different algorithms across various omics combinations. The log-rank test was used to assess statistical significance between survival curves. Abbreviations: CNV, copy number variation; Ex, gene expression; Me, DNA methylation.

**Supplementary Fig. 12.**
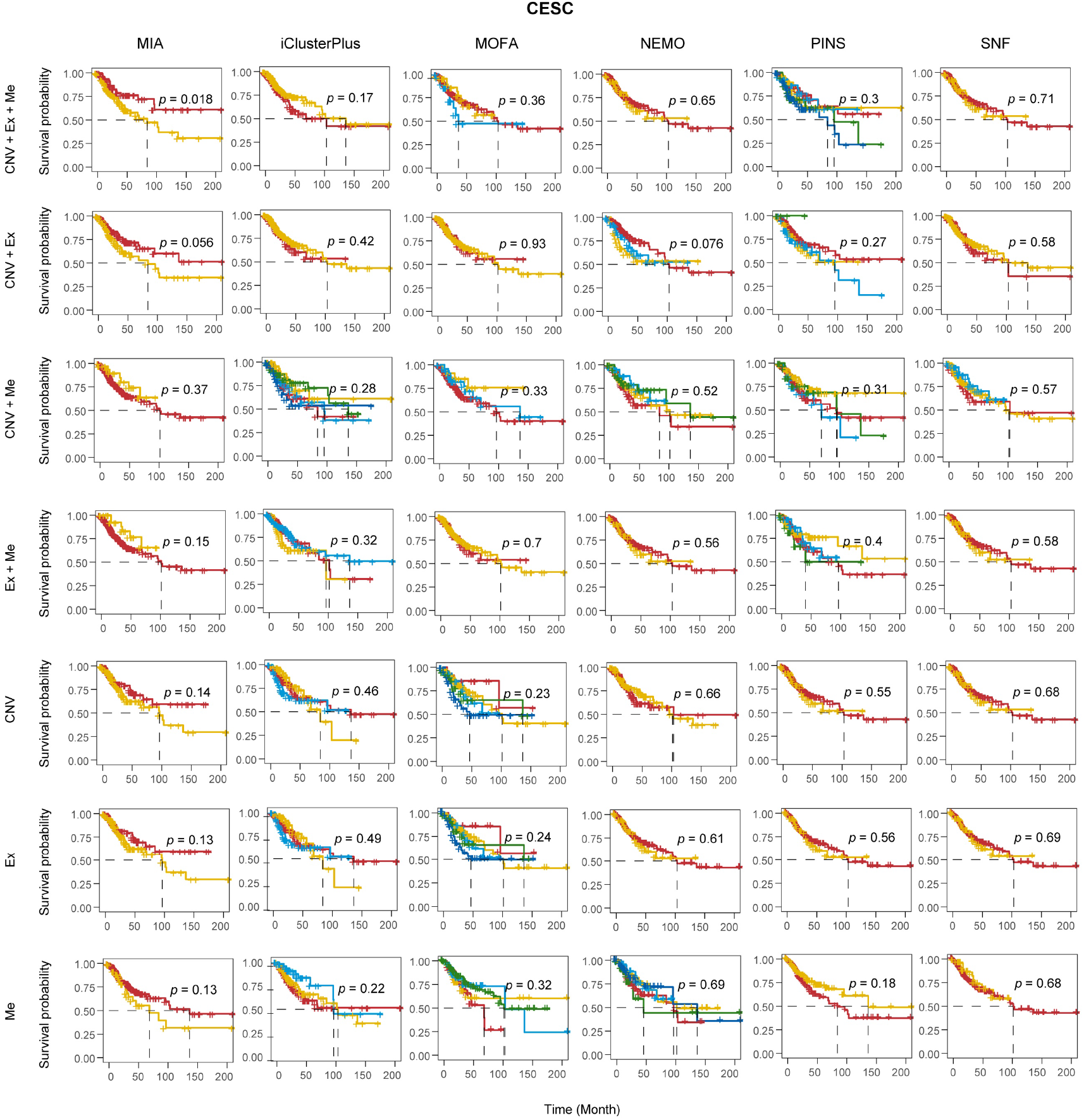
Kaplan-Meier survival analysis on TCGA CESC dataset clustered by different algorithms across various omics combinations. The log-rank test was used to assess statistical significance between survival curves. Abbreviations: CNV, copy number variation; Ex, gene expression; Me, DNA methylation.

**Supplementary Fig. 13.**
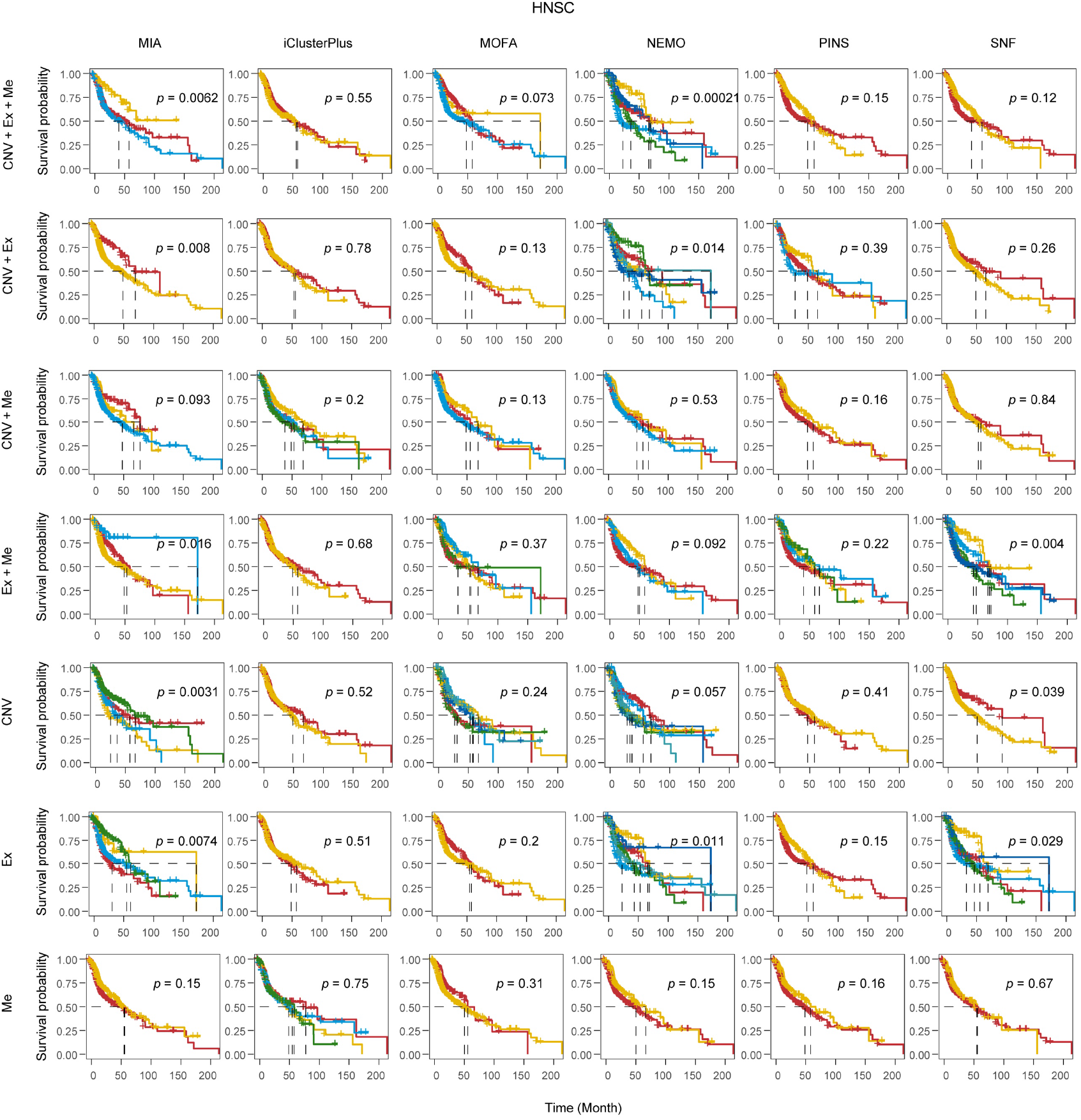
Kaplan-Meier survival analysis on TCGA HNSC dataset clustered by different algorithms across various omics combinations. The log-rank test was used to assess statistical significance between survival curves. Abbreviations: CNV, copy number variation; Ex, gene expression; Me, DNA methylation.

**Supplementary Fig. 14.**
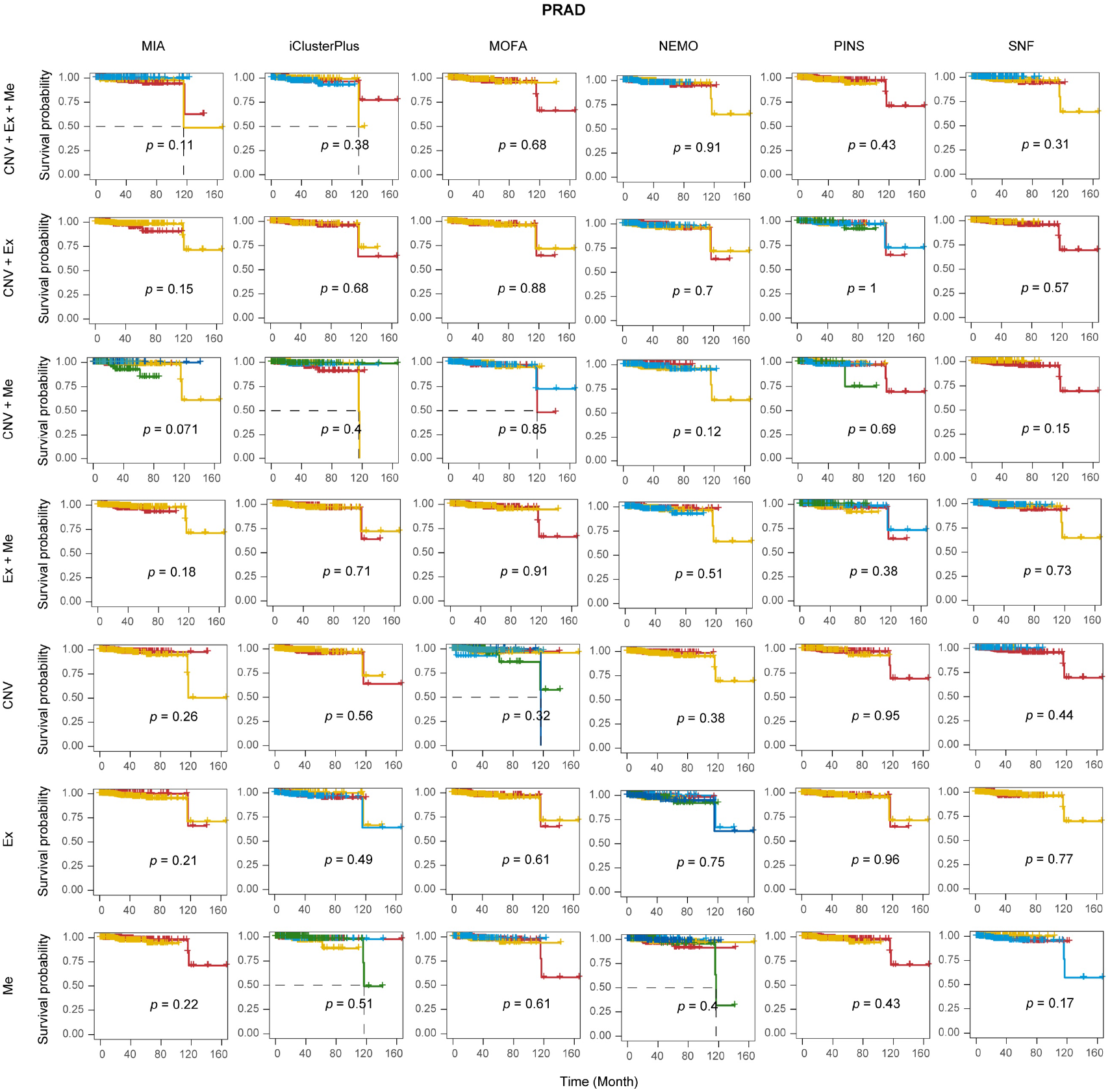
Kaplan-Meier survival analysis on TCGA PRAD dataset clustered by different algorithms across various omics combinations. The log-rank test was used to assess statistical significance between survival curves. Abbreviations: CNV, copy number variation; Ex, gene expression; Me, DNA methylation.

**Supplementary Fig. 15.**
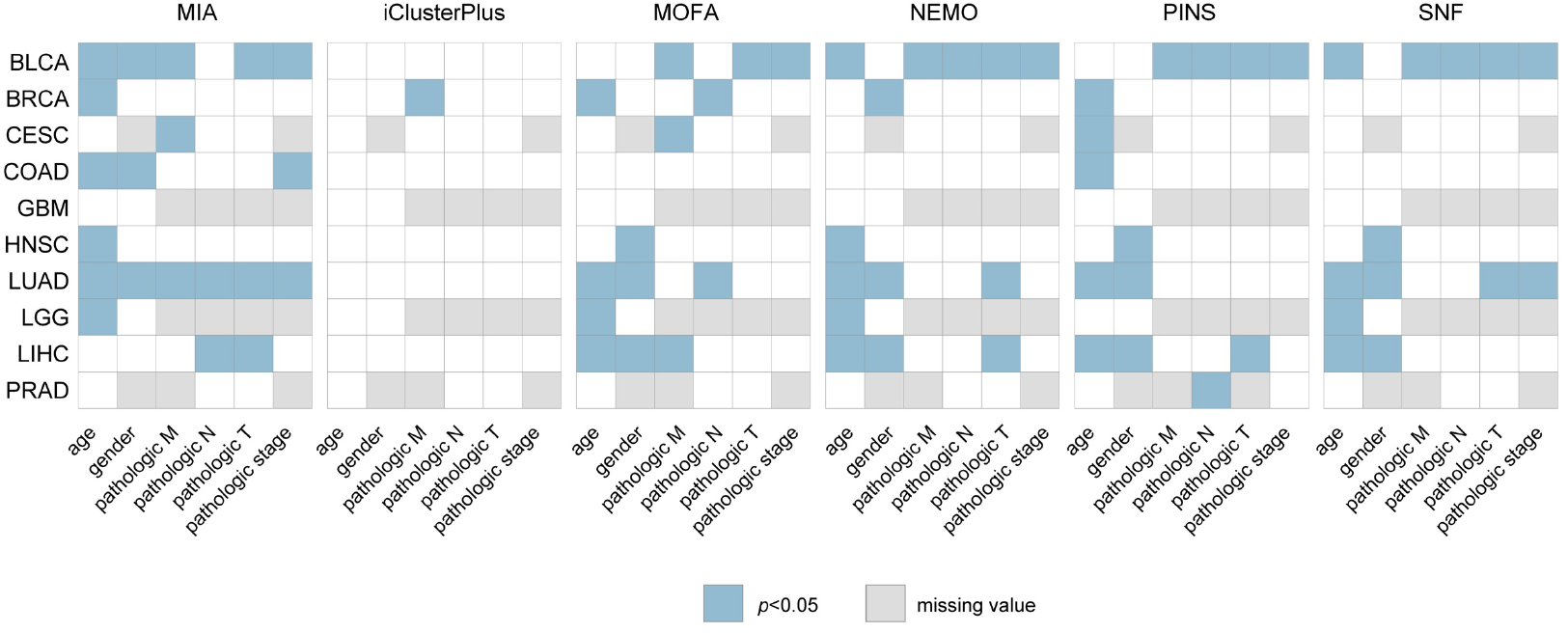
Enrichment of clinical records by different algorithms across ten cancer types. Clinical records include Pathologic M, metastasis status; Pathologic N, lymph node involvement; Pathologic T, tumor progression; Pathologic stage, overall pathologic stage.

**Supplementary Fig. 16.**
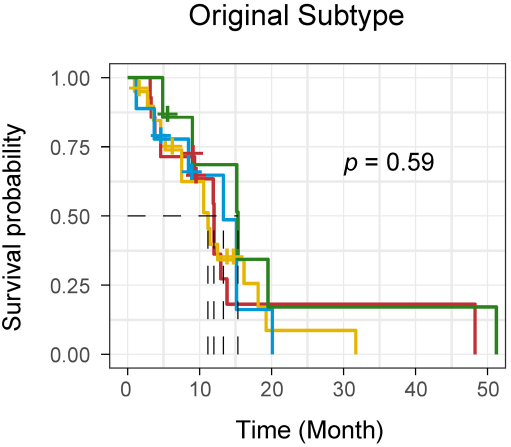
Kaplan-Meier survival analysis of GBM classical subtypes. The log-rank test was used to assess the statistical significance between survival curves.

**Supplementary Fig. 17.**
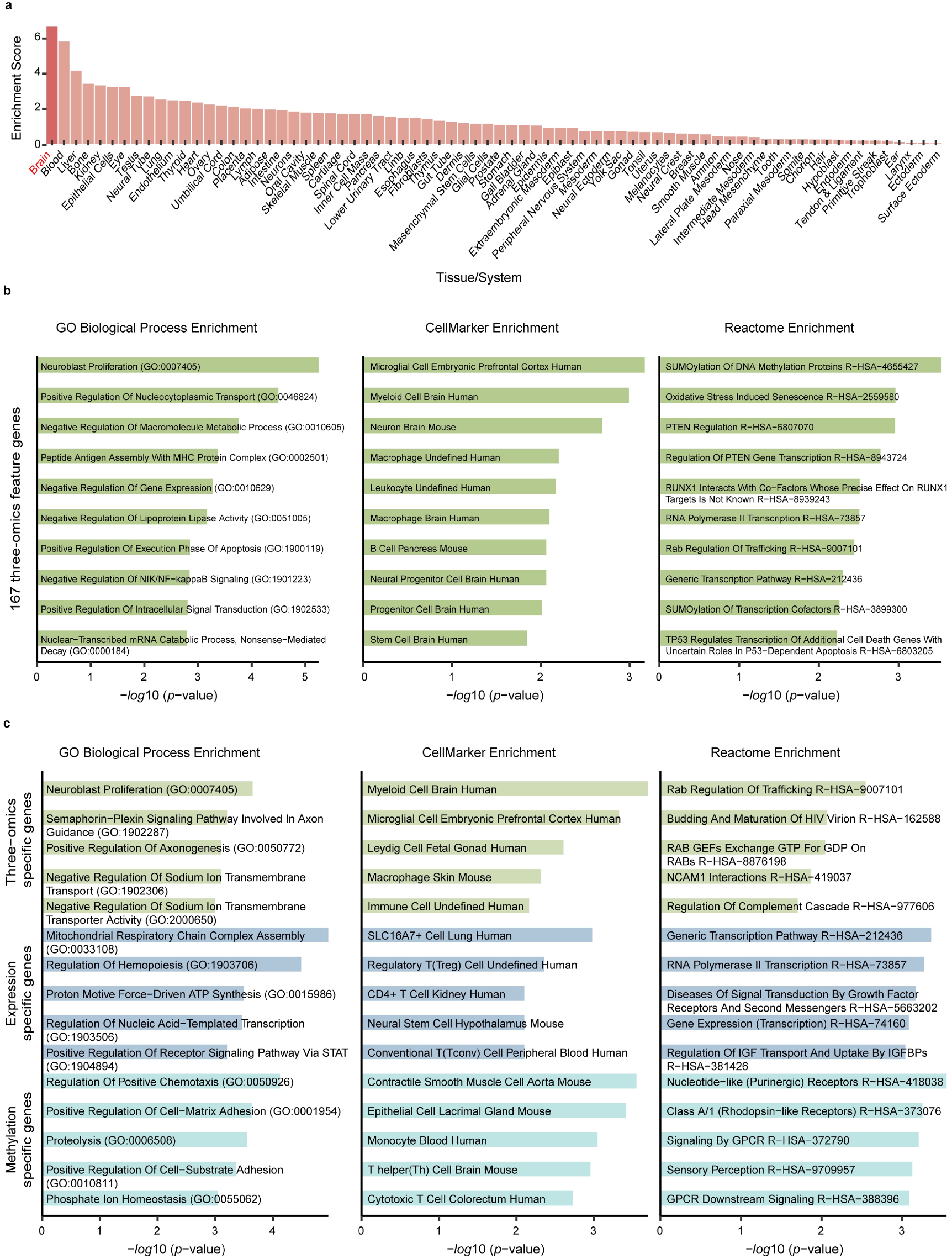
Functional enrichment of 167 GBM feature genes. (a) Enrichment scores of 167 feature genes across tissues/systems. (b) Enrichment analysis of 167 feature genes identified from multi-omics data. (c) Enrichment analysis of feature genes identified specifically from multi-omics, expression, and methylation data, respectively.

**Supplementary Fig. 18.**
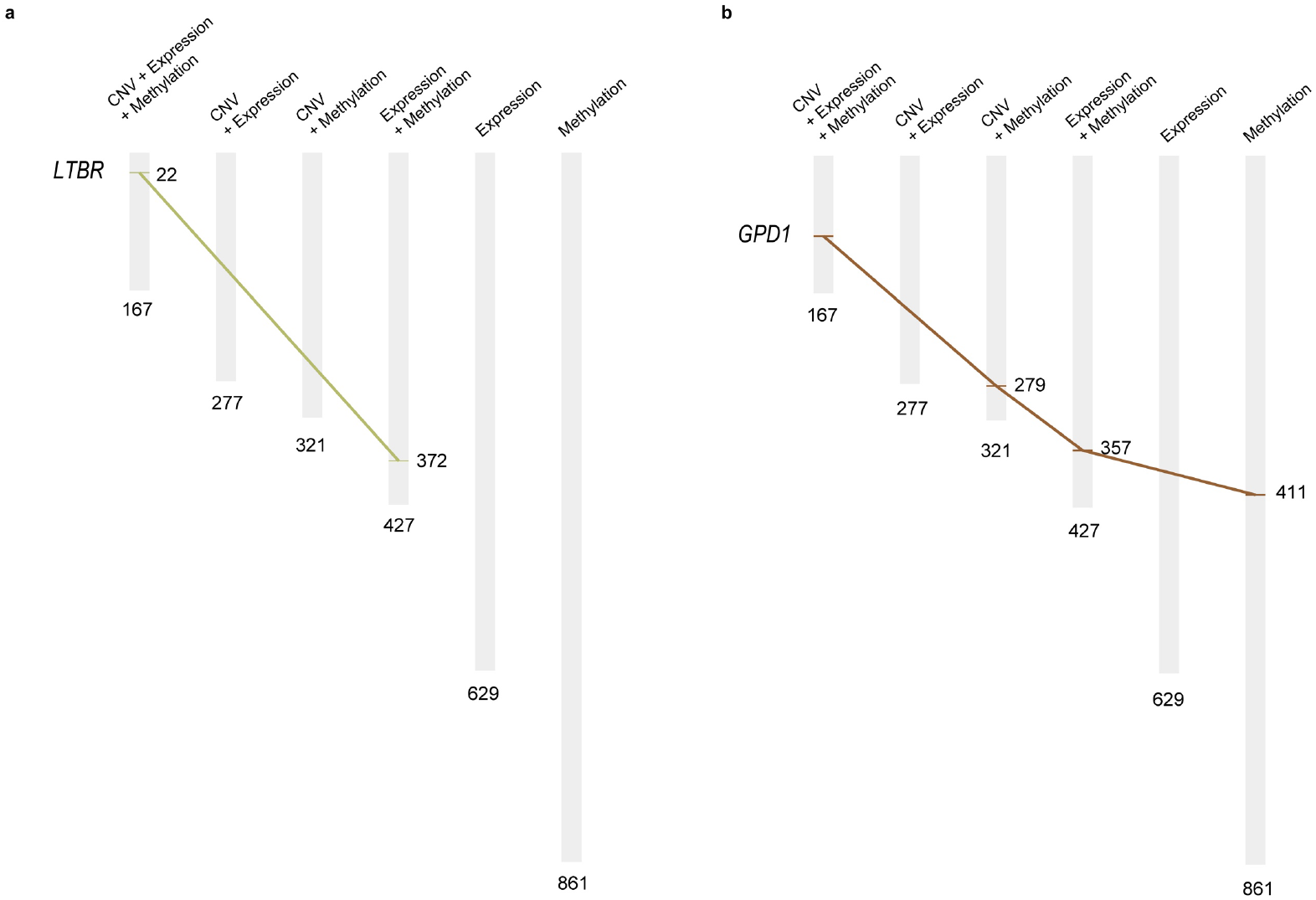
Importance ranking of *LTBR* (a) and *GPD1* (b) identified from different omics combinations. Feature genes identified by CNV (not shown) did not reach the threshold.

**Supplementary Fig. 19.**
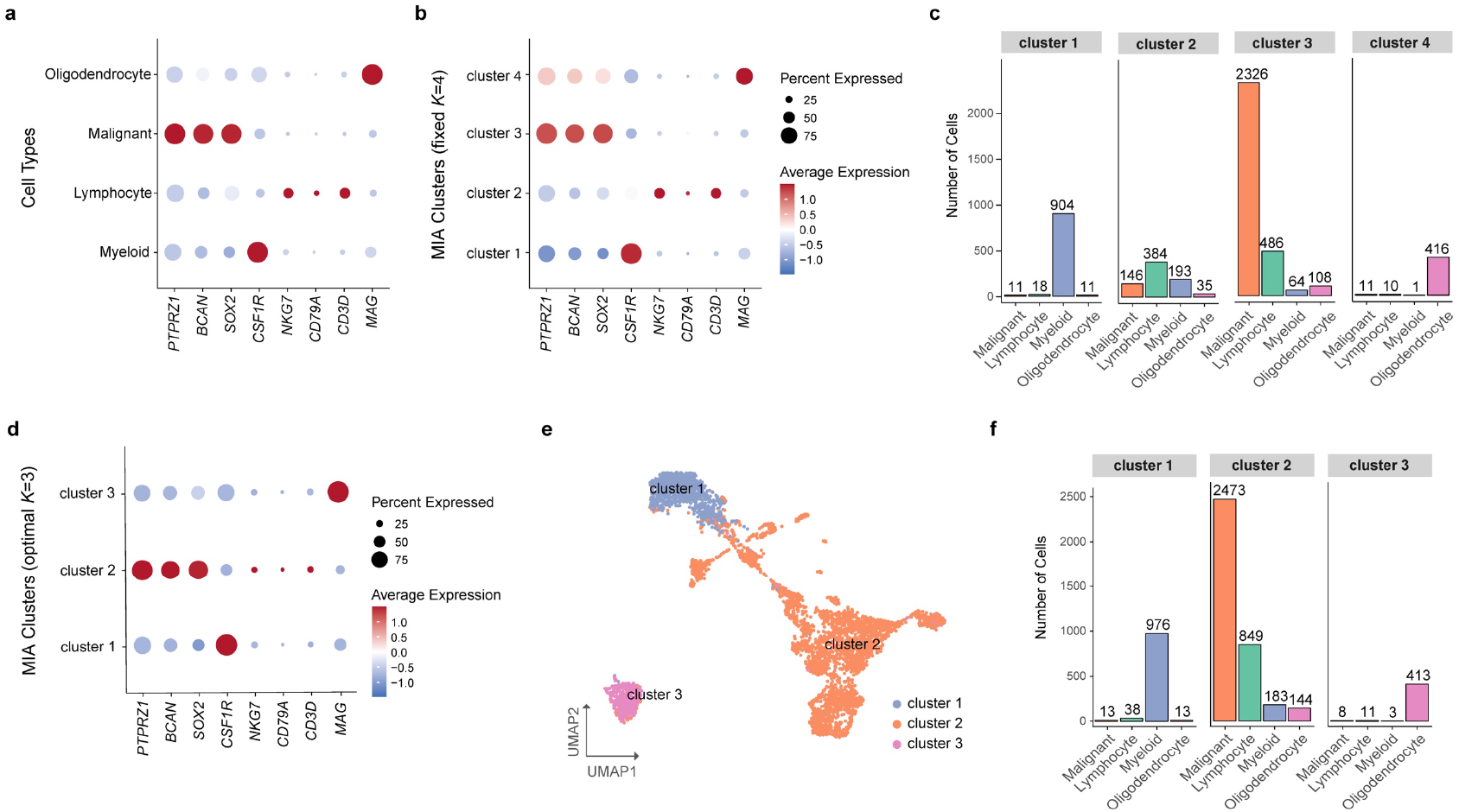
Gene expression on cell types. (a) Expression of cell markers across annotated cell types. (b) Expression of cell markers on MIA clustering with fixed *K*=4. (c) MIA clustering with the number of clusters fixed at *K*=4. (d) Expression of cell markers on MIA clustering with optimal *K*=3. (e) MIA clustering with the number of clusters automatically determined at *K*=3. (f) Proportional composition of annotated cell types within MIA clusters (*K*=4).

